# OpenAlea.HydroRoot: A modelling framework to dissect, predict and phenotype branched root hydraulic architecture

**DOI:** 10.64898/2026.03.19.713025

**Authors:** Fabrice Bauget, Adama Ndour, Yann Boursiac, Christophe Maurel, Laurent Laplaze, Mikael Lucas, Christophe Pradal

**Affiliations:** CIRAD, UMR AGAP institut, Montpellier, France; AGAP institut, Univ Montpellier,CIRAD, INRAE, Institut Agro, Montpellier, France; IPSiM, Univ Montpellier, CNRS, INRAE, Institut Agro, Montpellier, France; UMR DIADE, University of Montpellier, IRD (Research Institute for Development), Montpellier, France; Inria & LIRMM, Univ Montpellier, CNRS, Montpellier, France; International Maize and Wheat Improvement Center, Addis Ababa, Addis Ababa, Ethiopia

**Keywords:** OpenAlea, FSPM, root system architecture, hydraulic, plant modeling

## Abstract

Drought is a significant factor in agricultural losses, making it imperative to understand how root system architecture (RSA) adapts to environmental condition like water deficit. HydroRoot is a functional-structural plant model (FSPM) aimed at analyzing and simulating hydraulic and solute transport of RSA. The model integrates a static hydraulic solver, a coupled water-solute transport solver, a statistical generator of RSA based on Markov model, and a dynamic hydraulic model accounting for root growth. This paper presents the model, the mathematical description of the formalism of solvers, and use cases with their associated tutorials. Five use cases illustrate capabilities of HydroRoot, which has been successfully used for phenotyping root hydraulics across various species, including Arabidopsis, maize, and millet. The model-driven phenotyping method “cut and flow” is presented to characterize axial and radial conductivities on a given root genotype. Finally, three step-by-step tutorials provide a structured way to learn how to use HydroRoot 1) to simulate hydraulic on a given architecture, 2) to simulate water and solute transport on a maize root, and 3) to simulate hydraulic on two pearl millet genotypes with varying soil conditions. Hydroroot is an open-source package of the OpenAlea platform, with the code publicly available on Github. A comprehensive documentation is available with a reproducible gallery of examples.

## 1. Introduction

### 1.1. Root Hydraulic Architecture

Drought is one of the primary drivers of agricultural losses, accounting for nearly half of all agricultural disasters according to the latest FAO’s reports (doi:10.4060/cd7185en). Potential for drought tolerance in land plants arise from an interplay between their aerial part (stomatal conductance regulation, leaf adaptation, …) and their root system, which is responsible for water absorption from the soil. Thus, understanding how the hydraulic properties of Root System Architecture (RSA) evolve during growth, adapt to environmental condition such as soil water deficit and high-evaporative demand (Carminati *et al*., 2026), and vary naturally for a given species is of great importance. This is particularly true in the context of developing and selecting crop for improved drought tolerance. Functional– Structural Plant Models (FSPMs) of water transport at the RSA scale provide a robust framework for integrating architectural geometry, topology, and the spatial heterogeneity of transport parameters along the RSA (Passot *et al*., 2019). Hydraulic models of RSA have been developed for over two decades, beginning with the work of Doussan *et al*. (Doussan *et al*., 1998a,b), and further advanced by (Javaux *et al*., 2008; Draye *et al*., 2010; Couvreur *et al*., 2012; Zarebanadkouki *et al*., 2016; Meunier *et al*., 2017; Bouda *et al*., 2018).

Here, we present HydroRoot, a FSPM designed to estimate water and solute transport parameters and simulate RSA of cereals and Arabidopsis (Boursiac *et al*., 2022; Bauget *et al*., 2023; Rishmawi *et al*., 2023; Protto *et al*., 2024). It is a component of the OpenAlea platform (Pradal *et al*., 2008). HydroRoot represents root topology and geometry, including branching, root lengths and radii, transport properties, using a multiscale tree graph (MTG) (Godin & Caraglio, 1998; Pradal & Godin, 2020). The model integrates:

- a hydraulic model for static root structure,
- a coupled water-solute model for static structure,
- a module for RSA generation, and
- a hydraulic model for dynamic structure, accounting for growth.

### 1.2. Background to root hydraulic architecture modelling

Hydroroot has been used to phenotype the root hydraulics of different species: Arabidopsis (Boursiac *et al*., 2022), maize (Bauget *et al*., 2023; Rishmawi *et al*., 2023; Protto *et al*., 2024). The model was also used to simulate the development of observed pearl millet roots. HydroRoot is an original model in several aspects. The hydraulic module on static RSA uses the graph properties of MTGs to solve the system in an efficient way. The solute coupled to water associates radial and axial transport of solutes to classical water flow equations. Boursiac *et al*. (2022) proposed an original phenotyping protocol assisted by HydroRoot called cut and flow that allows to infer the axial and the radial conductivities of a given root from experiments.

## 2. Model description

### 2.1. Design of a root hydraulic model

The HydroRoot model is based on the Hydraulic Architecture of Root Systems, in other words, a transport model applied at the root architecture level (Doussan *et al*., 1998a).

Transport is modeled at tissue scale with two components: axial and radial. Axial parameters characterize the water flux along xylem vessels. Radial transport describes the flux of water and/or solute through cell layers surrounding the stele (epidermis, cortex and endodermis) and the vessels (xylem parenchyma). All peripheral layers are then modeled as a homogeneous medium. Transport parameters, as variables like solute concentration and pressures, are defined at any point along the RSA.

Root architecture leverages on MTG with vertices representing root segments. MTG allows accounting for complex topology of plant growth, for instance roots of different orders and different types (primary, seminal, crown, etc.). Each vertex has a cylindrical geometry characterized by a length *δl* (*m*) and a radius *r* (*m*) (Figure 1). Vertex indices are numbered in a prefix order, from parent to children direction, i.e., from base (index *i* = 0) to apex. In the following, each equation is written locally, at the scale of a root segment. All state variables, pressure and concentration, are associated either to xylem vessels or outside the root as boundary conditions. All parameters are defined at this scale. In the model implementation, variables and parameters are stored on the MTG as vertex properties.

**Figure 1:**
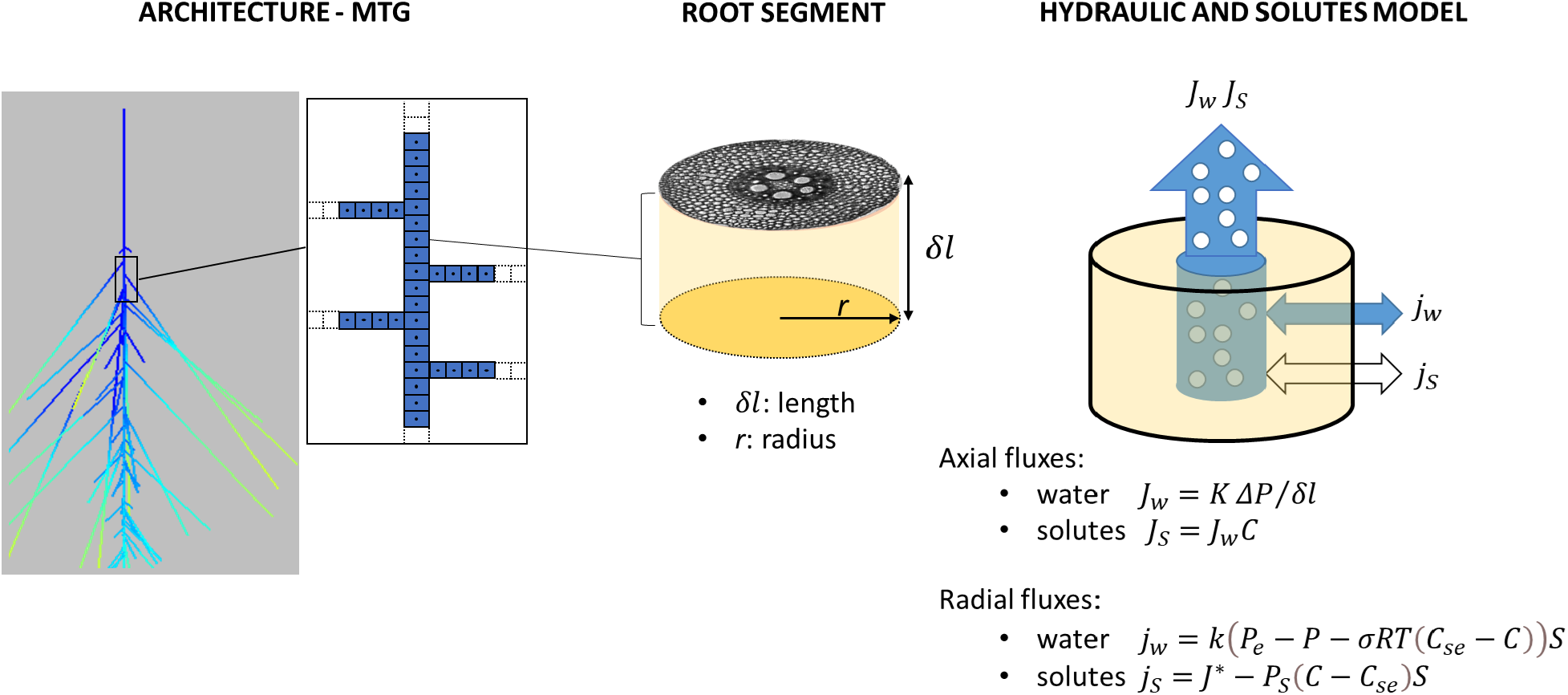
HydroRoot model: root is discretized as a MTG. Each MTG vertex represents a cylindrical root segment of radius r and length δl. Fluxes of water and solute are modeled on each root segment at axial (J_w_ and J_S_) and radial (j_w_ and j_S_) levels. K is the axial conductance, ΔP is the hydrostatic pressure difference along the root segment. C is the solute concentration in xylem vessels. k is the radial conductivity, P_e_ is the external hydrostatic pressure, P is the hydrostatic pressure inside the xylem vessels, σ is the reflection coefficient, R is the gas constant, T is the temperature, C_se_ is the external solute concentration, J^*^is the active uptake rate of solute, P_s_ is the radial permeability of the root peripheral tissues, and S is the area of the surface of exchange.

There are two hydraulic solvers: hydrostatic and hydraulic with solute transport. The hydrostatic model can be solved either directly using a MTG traversal, or using linear algebra. Solver of the solute model has been only implemented using a sparse matrix formalism. Both model types may have either homogeneous boundary condition (e.g. hydroponic condition), or heterogeneous boundary condition (modeling a heterogenous soil).

#### Major assumptions

- HydroRoot is soil less, i.e. soil is only viewed as a constant or variable boundary condition for state variables in 1D. There is no interaction between soil and root.
- Axial and radial hydraulic conductances depend only on the distance to the root tip, which serves as a proxy for age, at a given time.
- A root segment is considered homogeneous. There is no explicit description of the segment cell tissues. It is simply made of two concentric homogeneous compartments: 1) the stele with xylem vessels, and 2) the peripheral tissues.
- The model has been validated only on annual plants.

### 2.2. Root Architecture

In HydroRoot, the RSA data-structure can either come from phenotyping experiments (i.e. measurements or scans) via file importation, or generated using a simulation model that can produce a RSML output such as OpenSimRoot(Postma *et al*., 2017), CPlantBox(Schnepf *et al*., 2018), and OpenAlea(Pradal *et al*., 2008). Note that in previously published studies (Boursiac *et al*., 2022; Bauget *et al*., 2023; Rishmawi *et al*., 2023; Protto *et al*., 2024), HydroRoot was used to analyze and phenotype plants grown and measured in a hydroponic solution, i.e. under uniform external condition. Under such conditions, the distance to the base was sufficient to characterize the position of a given vertex.

#### Acquisition from Root Phenotyping

The Root System Markup Language (RSML) (Lobet *et al*., 2015) can be used as input in HydroRoot. RSML is a standard format that allows a complete three-dimensional description of the root architecture: root lengths, root angles, branching position. It is compatible with various root simulation software and phenotyping pipelines.

A simplified csv format can also be used in the specific case of cut and flow experiment where RSA is reconstructed from successively cut roots (Bauget *et al*., 2023). This file is composed of three mandatory columns: the lateral branching position on their parent, the lateral length and its order. A fourth optional column can represent the lateral diameter. With this format, the topology is preserved but not the geometry. This is not a problem in hydroponic condition, where the boundary conditions are constant. However, the geometry is needed when boundary conditions are variable to simulate a soil type.

#### Static Architecture Generation

HydroRoot provides a generator of RSA based on a stochastic process modeled by a first-order Markov chain on growing root axes (Lucas *et al*., 2008) using several parameters:

- the primary root length L;
- a nude tip length l_n_, that is, the root segment from the apex without any branching;
- an average internode length Δ;
- lateral length laws l_r_, which give lateral length according to position on parent axis on root order.

Two different laws are defined: one for first-order lateral root lengths and another for higher-order laterals. Boursiac *et al*. analyzed lateral root length as a function of branching position along the bearing axis, using 13 individual primary roots and 9 first-order laterals. They observed for both primary and lateral roots progressive and stochastic growth patterns, and proposed exponential (expovariate) fits to the experimental measurements as descriptive laws (Boursiac *et al*., 2022).

#### Dynamic Architecture Generation

A dynamic architecture can be generated by three types of methods: 1) RSA simulation, 2) 2D+t phenotyping, and 3) a hybrid method that simulates RSA growth and development from a static RSA observed at a final stage.

First, main RSA simulation software such as OpenSimRoot (Postma *et al*., 2017), CRootBox (Schnepf *et al*., 2018), or RhizoDep (Rees *et al*., 2025) provide a RSML format as output, that can be converted into a MTG. Second, some 2D+t Root phenotyping pipelines that capture RSA also export their results into RSML like RootSystemTracker (Fernandez et al., 2022), and RootNav 2.0 (Yasrab *et al*., 2019). Third, based on a static measured RSA that has been grown during T hours, we propose a simple model to add a time parameter (i.e. Days after Germination or DAG) to each vertex of the graph, given the following hypotheses:

- All the tips have the same age, i.e. they are growing.
- The growth of the longest root is uniform, i.e. the growth speed is constant.
- The delay before the emergence of a lateral is constant, and the growth speed is constant per root axis.

Based on these assumptions and with two parameters (age of the root and delay of lateral emergence), a date of appearance can be computed for each root segment.

### 2.3. Modeling axial fluxes

Locally, xylem vessels can be seen as pipes, in which sap flow can be modeled with a Hagen-Poiseuille’s law type. This law states that the incompressible flow of a viscous fluid in a circular pipe is proportional to the 4th power of radius times the hydrostatic pressure gradient. Having no knowledge of the shape of the xylem vessels, the law can be generalized as follows:

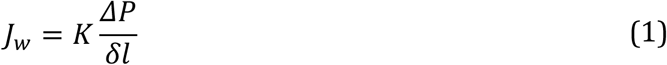

where *J*_*w*_ (*m*^3^.*s* ^−1^) is the axial water flux, *K* (*m*^4^. *s*^−1^. *MPa*^−1^) is the axial conductance, Δ*P* (*MPa*) is the hydrostatic pressure difference along the root element and *δl* (*m*) is the length of the element. The solutes inside the xylem vessels are transported by advection, therefore their axial flux *Js* (*mol*. *s*^−1^) is simply given by:

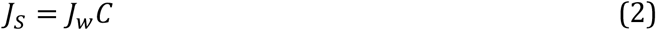

where *C* (*mol*. *m*^−3^) is the solute concentration in the sap.

### 2.4. Modeling radial fluxes

The cell layers surrounding xylem vessels are considered as an osmotic barrier through which water flux may be modeled with a Darcy’s law type with the flux being proportional to the water potential difference between the medium surrounding the root and the xylem vessels:

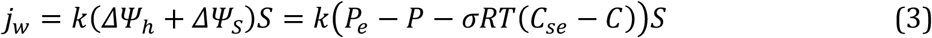

where *j*_*w*_ is the water radial flux, *k* (*m*. *s*^−1^. *MPa*^−1^) is the radial conductivity, Δ*Ψ*_*h*_ and Δ*Ψ*_*s*_ are the differences between the surrounding medium and the xylem vessels of hydrostatic water potential and of osmotic potential (*MPa*), *S* = 2*πrl* (*m*^2^) is the area of the surface of exchange, *P*_*e*_ is the external hydrostatic pressure, *P* is the hydrostatic pressure inside the xylem vessels, *σ* is the reflection coefficient, *R* is the gas constant (*J*. *K*^−1^. *mol*^−1^), *T* (*K*) is the temperature, *C*_*se*_ is the external solute concentration and C is the solute concentration in the xylem vessels.

The solute transport through the peripheral cell layers is described by a pump-leak model (Fiscus, 1975):

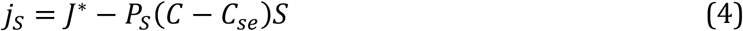

where *j*_*s*_ (*mol*. *s*^−1^) is the solute radial flux, *J*^*^ (*mol*. *m*^2^. *s*^−1^) is the solute active uptake rate, and *P*_*s*_ (*m*. *s*^−1^) is the radial permeability of the root peripheral tissues.

### 2.5. HydroRoot solvers

#### Hydrostatic solver: Efficient solving of Hydrostatic models on a tree graph

When the solute transport can be neglected, the governing equations (1) and (3) are reduced to *j*_*w*_ = *k*(*P*_*e*_ − *P*)*S*. By analogy with Ohm’s law, these equations can be written as follows (Boursiac *et al*., 2022):

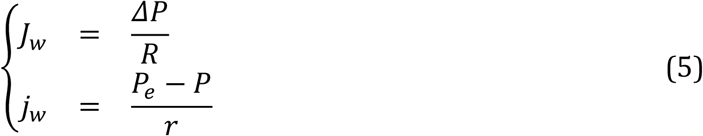

where axial and radial water flux, *J*_*w*_ and *j*_*w*_, may be seen as equivalent to electrical intensities, Δ*P* and (*P*_*e*_ − *P*) are analogous to potentials, and the inverse of conductivities are axial and radial resistances, *R* = δ*l*/*K* and *r* = 1/(*kS*). Therefore, a MTG is analogous to an electrical network and mathematical calculation can be done by a graph traversal Figure 2.

**Figure 2:**
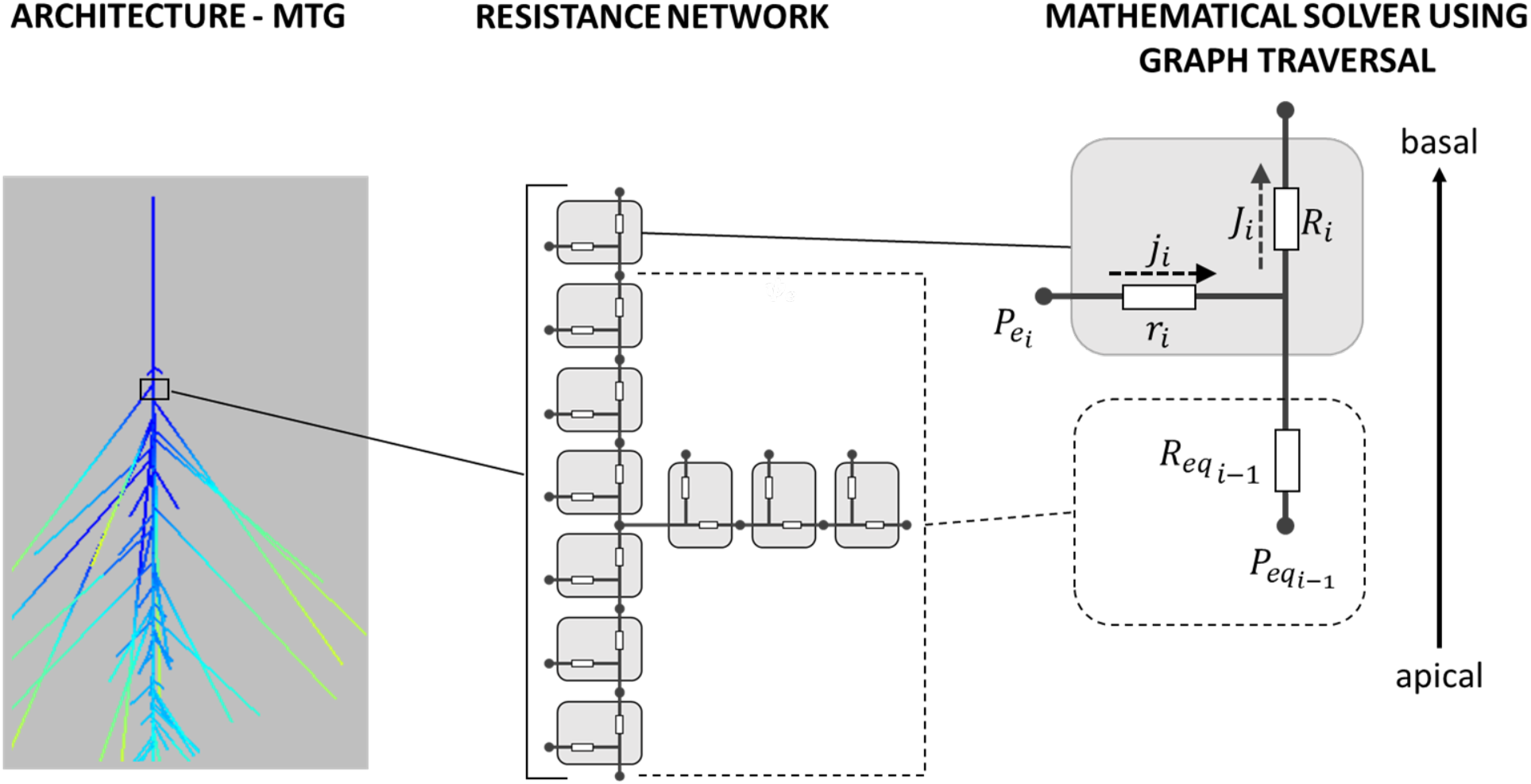
Hydrostatic solver implementation. RSA is modeled as a resistance network on which equivalent resistance can be calculated step by step from tips to base. 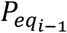 is the apical equivalent external hydrostatic pressure, 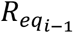 is the apical equivalent resistance, 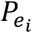 is the external hydrostatic pressure at current element i, r_i_ is the radial resistance of the current element i, R_i_ is its axial resistance, j_i_ and J_i_ are radial and axial fluxes respectively.

According to equation (5), each element can be seen as two linear dipoles in parallel. According to Thévenin’s theorem each element can be represented by an equivalent dipole with a single potential and a single equivalent resistance (Prusinkiewicz *et al*., 2007). Each element being in series with the previous equivalent dipole, we are able to calculate the equivalent resistance 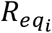 from the tip to the element *i* as follows:

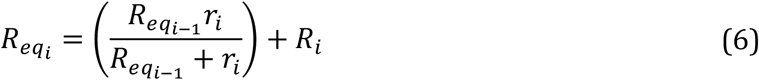

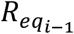 is the apical equivalent resistance, *R*_*i*_ is the axial resistance of the current element and *r*_*i*_ is its radial resistance. Then, by proceeding element by element from each tip, and because branching roots are analogous to a parallel network, we can calculate the equivalent resistance of the entire root and consequently its equivalent hydraulic conductance *K*_*eq*_.

The basal outgoing flux (*J*_*v*_) from the root can then be calculated according to the boundary conditions. For example, with a uniform water pressure surrounding the root, which is the case of hydroponic experiments (Boursiac *et al*., 2022), *J*_*v*_ is given by:

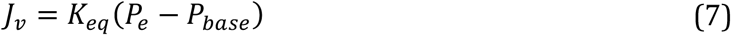

with *P*_*e*_ and *P*_*base*_ being the external pressure and the basal pressure, respectively.

With a non-uniform boundary condition, typically within a soil, an equivalent pressure must be calculated step by step starting from the tip with the following equation:

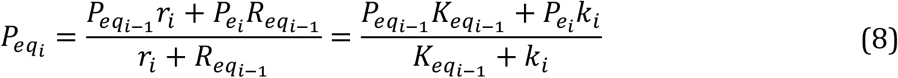

And finally, with *P*_*eq*_ being the equivalent external pressure after calculation over the whole root, *J*_*v*_ is given by:

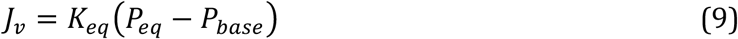

#### Solute solver: Generic solving all models using sparse matrix

With solute transport, there are two more equations coupled with previous ones making the step-by-step electrical network solver no longer valid. The root is then seen as a continuous medium with the system of equations discretized by finite differences and the system solved by linear algebra (see Figure 3).

**Figure 3:**
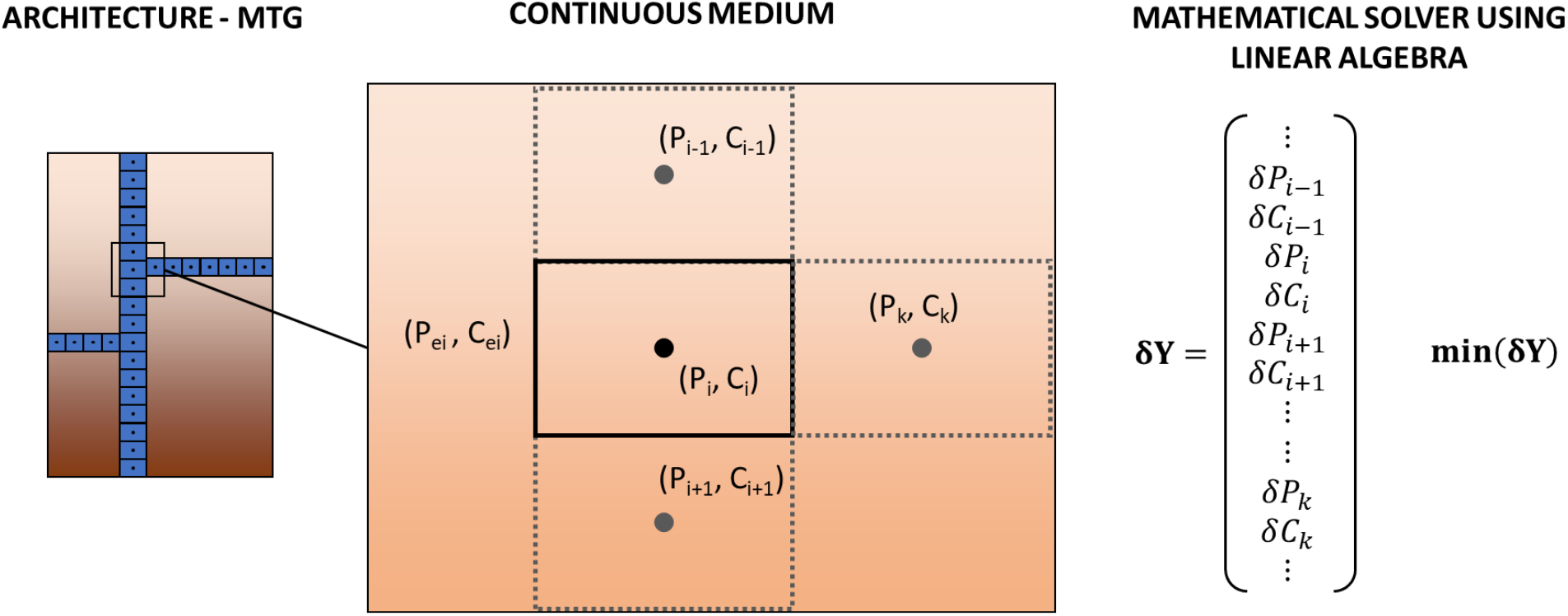
Solute solver implementation. The RSA is a continuous medium discretized with mesh size corresponding to vertices length. The principle is to minimize vector **δY** with a Newton-Raphson scheme.

The system of equations is obtained by applying the mass balance equation locally. In other words, at any point of the root, the outgoing flux of a root segment equals all ingoing fluxes, and this is true for water and solutes. Using equations (1) to (4), the system of equations governing water and solute transport in the root is:

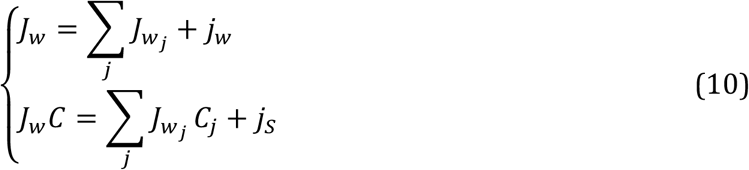

The former equation is the equation for water transport, where *J*_*w*_, the axial outgoing flux, equals the sum of the axial fluxes of any branched root segment plus the radial flow of water entering the root element. The latter equation governs solute transport where the axial outgoing solute flow by advection equals the fluxes coming from any branched roots and the incoming solute radial mass flow.

In the following, the subscript *w* will be dropped, *J* will always refer to axial water flux. The MTG structure is used as a discretization pattern where vertices correspond to elementary root segments. Vertices are numbered from the root base to the apex. Each vertex has properties length *δl* and radius *r, δl* is then used as mesh spacing. Considering vertex *i* the system of equations 10 is discretized as follows:

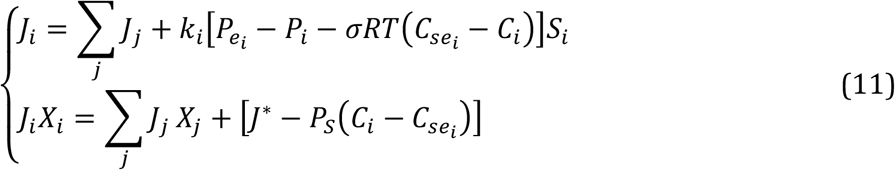

*X*_*i*_ represents concentration of solute according to flow direction between element *i* − 1 and *i*. Subscript *j* refers to the children of root segment *i*.

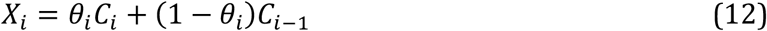

with *θ*_*i*_ = 0 if *P*_*i*−1_ > *P*_*i*_ and *θ*_*i*_ = 1 otherwise. This is an upwind scheme based on upstream weighting of flux that is commonly used in porous media modeling for stability reason (Aziz & Settari, 1979). *X*_*j*_ is the concentration of solute flowing between vertex *i* and its children. *J*_*i*_ and *J*_*j*_ are the axial flow from the element *i* to its parent *i* − 1 and the axial flow from the children *j* to *i*, respectively:

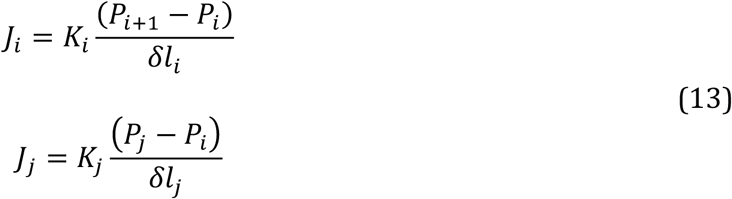

Then system Eq. 11 can be rewritten at the vertex *i* as follows:

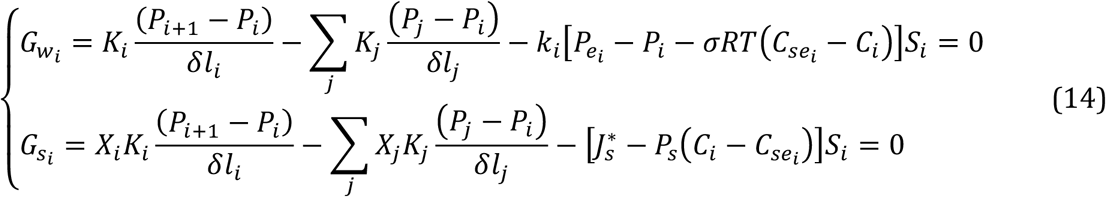

These equations can be seen as components of a vector **G**. This is now a minimization problem **G** = 0 with two unknown vectors **P** and **C**, the hydrostatic pressure and concentration of solutes, respectively, over all the root. In addition to this system, there are the following boundary conditions:

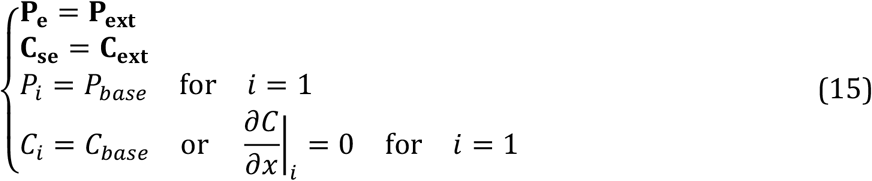

where **P**_**ext**_ and **C**_**ext**_ are the vectors of external hydrostatic pressure and concentration, all around the root, *P*_*base*_ and *C*_*base*_ are the hydrostatic pressure and concentration at the base of the root. The system is solved using a Newton-Raphson’s method that consists of solving iteratively:

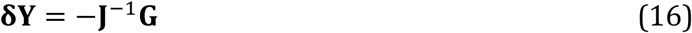

until **δY** is close to a null vector according to a convergence criterion. **δY** is the vector with the incremental difference on *P* and *C*:

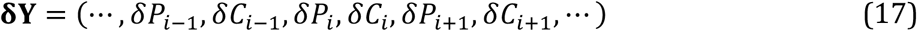

**J** is the Jacobian of **G** according to *P* and *C*:

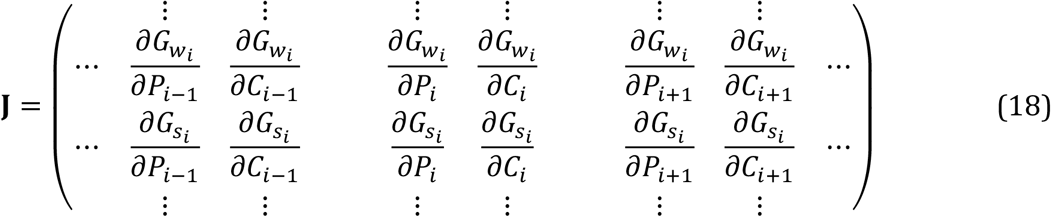

As shown by the system of equations (14), 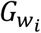 and 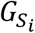 do not depend only on *P*_*i*_ and *C*_*i*_, but may also depend on variables in neighbors *j*. Therefore, **J** is not a band matrix but a sparse matrix. The inversion of **J** is done by LU decomposition using the sparse.linalg.splu from the SciPy Python library (Virtanen *et al*., 2020).

#### Boundary conditions

The hydrostatic pressure and concentration outside the root, **P**_**ext**_ and **C**_**ext**_ are local boundary conditions. Therefore, these components are at the root segment scale and may vary according to boundary conditions. For instance, homogeneous conditions lead to uniform *P*_*e*_ and *C*_*se*_ values along the MTG, and heterogeneous conditions correspond to different values associated with root segment positions. Thus, a soil with properties varying with depth can be modeled by associating segment at a depth *z* with *P*_*ext*_ (*z*) and *C*_*ext*_ (*z*).

### 2.6. Cut & Flow: Model inversion

Measuring axial and radial conductance is challenging. In pressure chambers, we propose to phenotype the hydraulic architecture using the hydrostatic solver of HydroRoot with a method called Cut and Flow (CnF, Boursiac *et al*., 2022). It consists of measuring efflux *J*_._ of an RSA in a pressure chamber on a root successively cut at decreasing distances from the root base:

1. Efflux is measured on the complete RSA.
2. Root is cut a first time at a certain distance from the root base. The efflux is measured on the remaining root in the pressure chambre.
3. Root is cut a second time and the efflux measured.
4. And so on as many times we want. Usually between 3 to 6 cuts are done.

The experimental data is then a flux versus a distance to root base. Finally, the CnF method of HydroRoot is used to find the unknown *K* and *k* that allow to fit the data.

This method has been extended to the root of maize under water stress by adding solutes and osmotic processes (Bauget *et al*., 2023; Protto *et al*., 2024). In addition to the CnF data, *J*_*v*_(*P*) data of efflux measured on the entire root at different pressures were added. Thus, using the CnF method of HydroRoot specific to solute solver *K, k, J*^*^ and *P*_*s*_ are then determined by fitting both data.

The main steps of the fitting process by reverse modeling are:

- setting the parameter values as first guesses
- minimizing *F* using optimize.minimize function of the SciPy Python library (Virtanen *et al*., 2020)

### 2.7. Non-permeating solutes

In the solute solver, the presence of non-permeating solutes can be accounted for. This possibility was implemented to be able to analyze CnF experiments with high molecular weight polyethylene glycol (PEG) in the hydroponic solution, which cannot cross plant tissues (Bauget *et al*., 2023). This addition does not fundamentally modify the general model description. However, it obviously changes external osmotic potential and, more importantly, when roots are cut, PEG molecules enter the xylem vessels, thereby changing both the viscosity and the osmotic potential within the xylem. Further details are provided in Bauget et al. (2023) and in the documentation.

## 3. Use Cases

### 3.1. Determining the role of RSA, axial and radial conductivities in *Arabidopsis thaliana*

CnF have been used to investigate the interplay between radial and axial conductivities of adult Arabidopsis Columbia-0 (Col-0) plants grown hydroponically (Boursiac *et al*., 2022). The plants were 21 days old and presented a complex and highly branched RSA. By image analysis of root cross sections, the authors observed that the root radius was homogeneous within each root order and was reduced by 40% between two successive orders. From these cross sections, the axial conductance was calculated from the radii of xylem vessels using the Hagen-Poiseuille’s law. Once *K* was determined, the radial conductivity was estimated using HydroRoot model on ten individual plants.

### 3.2. Phenotyping root hydraulic architecture using cut & flow experiments

Boursiac et al. also concomitantly determined axial and radial conductivities by fitting CnF data via model inversion (see paragraph 2.6). They showed, as previously reported (Tixier *et al*., 2013; Barry *et al*., 2025), that Hagen-Poiseuille’s law overestimated *K* by a factor of 4 to 6. They validated the model on Arabidopsis *eskimo1* mutants, which exhibit defects in xylem vessels. CnF analysis revealed a significant decrease in axial conductivity (by up to 19-fold) as well as a reduction of *k* by up to 55%.

### 3.3. Hydraulic responses to water deficit of maize roots

HydroRoot was employed to study the hydraulic properties of 11-day-old maize roots grown hydroponically under standard and water deficit (WD) conditions (Bauget *et al*., 2023; Protto *et al*., 2024). Bauget *et al*. showed that under WD conditions the osmotic component of root transport cannot be neglected. The authors therefore added a solute transport component (eq. 2 and 4) to the hydrostatic model (eq. 1 and 3). Using CnF experiments analyzed with HydroRoot, Protto *et al*. demonstrated first that the primary root exhibited a somewhat higher hydraulic conductivity than seminal roots. In addition, a WD reduced these parameters in both root types, together with root growth. In both root types, the profile of axial conductance was shifted to the root tip. Thus, the overall root hydraulic architecture was dramatically affected by the WD. Yet, the effects of WD on axial conductance could simply be explained by the reduction in root growth rate, assuming that the developmental process of xylem differentiation was not altered by the WD.

### 3.4. Characterizing natural hydraulic variation of maize RSA

Rishmawi *et al*. (2023) investigated the genetic variation of root hydraulic architecture within a panel of 224 maize inbred Dent lines (Rishmawi *et al*., 2023). They defined a core genotype subset of 13 lines to characterize in more details the architectural, anatomical, and hydraulic traits in the primary (PR) and seminal roots (SR) of hydroponically grown seedlings. They reported substantial genotypic variability in root hydraulic conductivity (*L*p_r_), PR size, lateral root size, as well as in the size and number of late metaxylem vessels. Later traits were positively correlated with *L*p_r_. Inverse modeling using HydroRoot further revealed pronounced differences among genotypes in xylem conductance profiles.

### 3.5. Genetic variation of pearl millet hydraulic architecture

Pearl millet is a major crop in Africa and India, reputed for being a drought tolerant crop (Shrestha *et al*., 2023). Passot et al. investigated the root anatomy and root architecture of Pearl Millet to decipher the underground mechanisms of its drought tolerance (Passot *et al*., 2016). They studied a panel of inbred lines for their natural variability in root architecture, and selected two highly contrasted lines (line ICMB 98222 and LCICMB1 – respectively lines 57 and 109 in the panel) to be further investigated for their hydraulic behavior (Passot, 2016) through pressure chamber measurements and inverse modeling of their root hydraulic architecture using HydroRoot. This particular use case of HydroRoot differs from the Maize and *Arabidopsis* use cases as the simulations are done on a growing architecture (using explicit growth time) and with a limited 1D environment representation (vertical gradient of external hydrostatic pressure).

## 4. Tutorial

### 4.1. Workflow

All tutorials follow the same workflow steps:

1. Acquisition or simulation of RSA.
2. Setting axial and radial parameters and boundary conditions.
3. Solving hydraulic architecture with dedicated solver.
4. Visualization and analysis.

In this tutorial section will show three examples: Arabidopsis in hydroponic condition, maize with solute transport and millet in a soil. These tutorials can be found as Jupyter notebook on the HydroRoot Github repository : https://github.com/openalea/hydroroot.

### 4.2. Quick start

*HydroRoot* is implemented in Python, leveraging the Multiscale Tree Graph (MTG), (Pradal & Godin, 2020) from the OpenAlea platform ((Pradal *et al*., 2008; 2015), https://openalea.readthedocs.io). *HydroRoot* is publicly available under the open-source license CecILL-C.

It can be installed on multiple platforms such as Linux, macOS, and Windows. Installation requires the package manager Conda (https://www.anaconda.com/docs/getting-started/miniconda/main). We recommend to install it via https://github.com/conda-forge/miniforge. Then, HydroRoot can be installed as User or Developer.

#### User installation

In a terminal, or in “miniforge prompt” for Windows users, just run the command:

~~~
              mamba create -n hydroroot -c conda-forge -c openalea3
              openalea.hydroroot jupyterlab
~~~

This will create a conda environement called hydroroot with openalea.hydroroot and all its dependencies installed. Jupyterlab will be also installed allowing to play the different notebooks present in the repository.

#### Developer installation

It is needed to install Git (https://git-scm.com/). Then to clone the project repository by running in a terminal the following command:

~~~
              git clone https://github.com/openalea/hydroroot.git
~~~

This will create in the current directory a directory “hydroroot”. Running the following:

~~~
              cd hydroroot
              mamba env create -f conda/environment.yml
~~~

This will create a conda environment, called `hydroroot_dev`, with all HydroRoot dependencies. HydroRoots will be locally installed from the source code as an editable install, using the command ‘pip install -e .’. To run the available notebooks, Jupyterlab must be installed in the environment “hydroroot_dev”.

A detailed documentation can be found at https://hydroroot.readthedocs.io. This documentation presents several Jupyter notebook examples that are available also in the github repository under the directory ‘doc/example’. They can be played interactively on the cloud using binder, by clicking on the binder badge in README page.

### 4.3. Model parameters

The main model parameters are summarized in Table 1:

**Table 1:** Parameters of HydroRoot model.

| Symbol | Parameter | Type | Range or value | Unit |
| --- | --- | --- | --- | --- |
| **Architecture parameters** |  |  |  |  |
| $r_{\text{ref}}$ | Reference radius<br>(radius of the primary root) | float | $10^{-5}$ to $10^{-3}$ | m |
| $\beta$ | Radii decrease factor<br>(between successive order) | float | 0.3 to 0.7 | m/m |
| $\delta l$ | Root segment length | float | $10^{-3}$ to $10^{-4}$ | m |
| **Additional architecture parameters for the generator** |  |  |  |  |
| L | Primary root length | float | 0.04 to 0.50 | m |
| $l_r$ | Laterals length laws<br>(vs relative distance to tip) | float | 0.00 to 0.12 | m |
| $\Delta$ | Average internode length | float | $10^{-3}$ to $3 \cdot 10^{-3}$ | m |
| n | Maximum order | integer | 4 | - |
| $\alpha$ | Branching variability | float | 0.25 | - |
| $l_n$ | Nude length | float | 0.02 | m |
| **Hydraulic parameters** |  |  |  |  |
| $k$ | Radial conductivity | float | $10^{-8}$ - $10^{-6}$ | $\text{m}\cdot\text{s}^{-1}\cdot\text{MPa}^{-1}$ |
| $K$ | Axial conductance<br>(vs distance to tip) | float | $10^{-13}$ to $10^{-10}$ | $\text{m}^4\cdot\text{s}^{-1}\cdot\text{MPa}^{-1}$ |
| **Experimental parameters** |  |  |  |  |
| $J_v$ | Water flux<br>(input for the K and k adjustment) | float | $10^{-3} - 1.0$ | $\mu\text{l.s}^{-1}$ |
| $P_e$ | External hydrostatic pressure | float | 0.4 | MPa |
| $P_{base}$ | Hypocotyl hydrostatic pressure | float | 0.1 | MPa |
| **Solute parameters** |  |  |  |  |
| $C_{se}$ | External concentration of permeable solutes (i.e. solutes able to cross plant tissues) | float | 13.96 | $\text{mol.m}^{-3}$ |
| $\sigma$ | Reflection coefficient | float | 0 to 1 | - |
| $J^*$ | Solute active uptake rate | float | $10^{-7}$ | $\text{mol.m}^2.\text{s}^{-1}$ |
| $P_S$ | Radial permeability | float | $10^{-9}$ | $\text{m.s}^{-1}$ |

**Table 2:** Pearl millet architectural parameters (Passot et al., 2016; Passot, 2016).

| Inbred line | Number of plants | Mean (± std) Primary Root Length (cm) | Mean number Crown Roots (± std) | Mean number Lateral roots (± std) |
| --- | --- | --- | --- | --- |
| 57 | 46 | 10.3 (6.4) | 0.17 (0.68) | 3.1 (3.9) |
| 109 | 76 | 20.2 (8.6) | 0.54 (1.1) | 14.1 (14.3) |

### 4.4. Tutorial 1: simulate hydraulic on a given architecture

The purpose of this tutorial is first to run simulations of an Arabipdopsis root grown in hydroponic conditions and measured using pressure chambers. The second purpose is to highlight how changes in radial and/or axial conductance impact water uptake. Hydroponic conditions are characterized by a homogeneous external hydrostatic pressure for which details on the hydrostatic solver can be found in Boursiac et al. 2022.

#### Module import

After HydroRoot installation, launch an ipython session. First import needed Python modules:

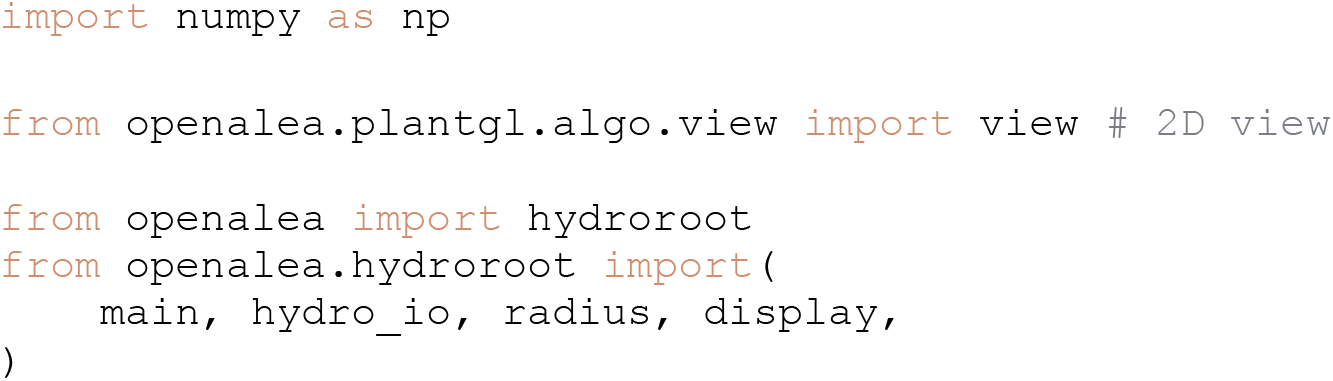

#### RSML to MTG

Read the RSML file and convert it to a continuous MTG. This file is located on https://github.com/openalea/hydroroot/blob/main/tutorial/data/arabidopsis.rsml

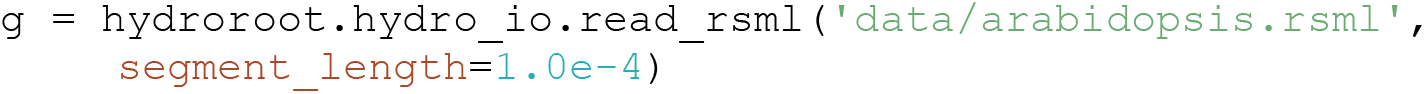

#### Set the root radii

The radius setting follows the Arabidopsis model described in Boursiac et al. 2022. The root radius is set according to the radius of the primary root *r*_*ref*_ = 7 10^−5^*m* and the order of the lateral as follows: *r*_*lat*_ = *β*^*order*^*r*_*ref*_. *β* = 0.7 is the radius decreasing factor between root orders.

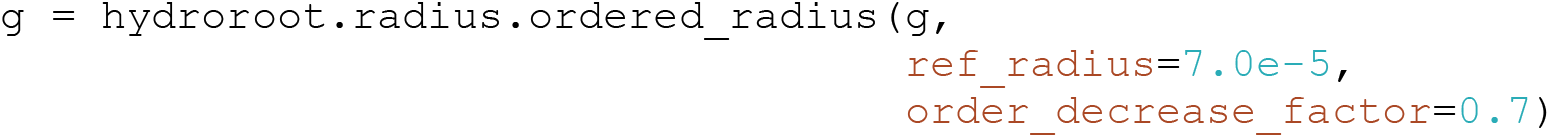

Compute the position of each segment relative to the axis bearing it

~~~
              g = hydroroot.radius.compute_relative_position(g)
~~~

#### Set hydraulic properties

We set the radial and axial conductivities as profiles, i.e. the property versus the distance to root tip in *m*. The radial conductivity k is in 10^−9^ *m. s*^−1^. *MPa*^−1^ and the axial conductance K is in 10^−9^ *m*^4^. *s*^−1^. *MPa*^−1^

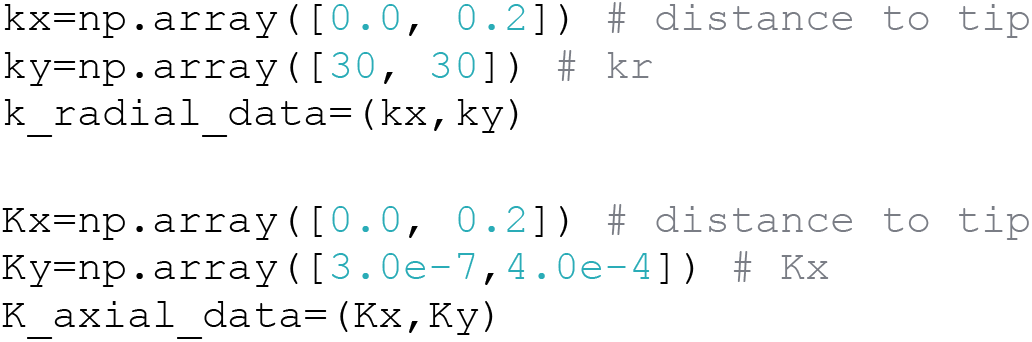

#### Flux and equivalent conductance calculation

The simulation is done for a detopped root in a hydroponic medium at 0.4 MPa with its base at 0.1 MPa.

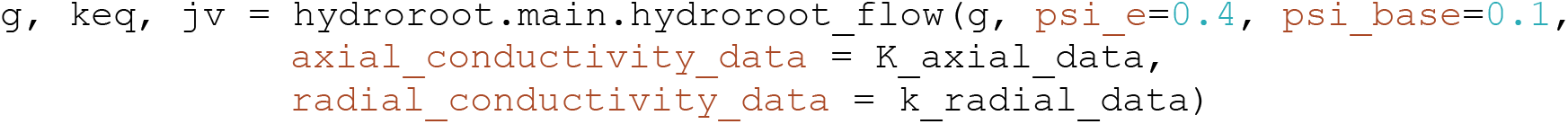

Expected results are a sap flux of 2.88 10^−3^ μL. S^−1^ and an equivalent root conductance of 9.60 10^−12^ *m*^3^. *s*^−1^. *MPa*^−1^

#### Visualisation

We can plot heatmap of different variables on the root architecture in 3D or 2D. First, we need to build a scene from the MTG with the proper variable to display. Note that variables are saved as MTG properties. For instance, if we want to display in 2D the water uptake in *μL. s*^−1^, i.e. radial incoming water flux, on the root with its scale bar (Figure 4A), we run the following commands:

~~~
          s = hydroroot.display.mtg_scene(g, prop_cmap=‘j’)
          view(s)
          hydroroot.display.property_scale_bar(g, ‘j’)
~~~

**Figure 4:**
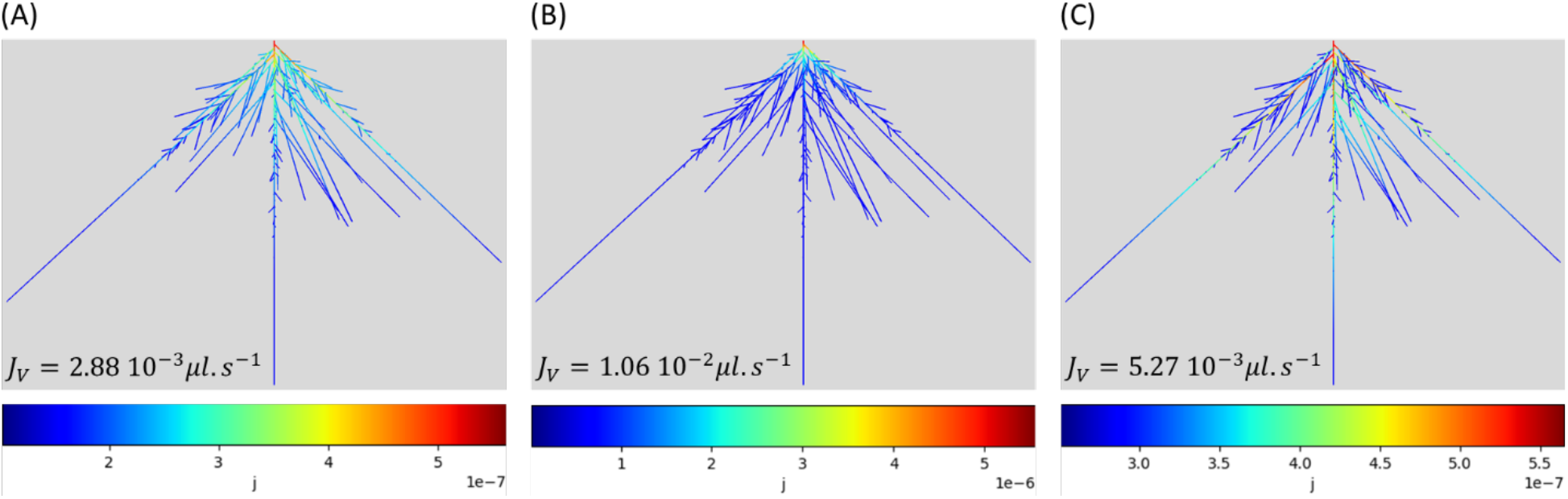
heatmap of water uptake of an Arabidopsis root with three different sets of conductivities with k uniform and K as a profile. (A) k_A_=30, K_A_ (3 10^-7^, 4 10^-4^) vs length (0, 0.2), (B) k_B_=10k_A_, K_B_=K_A_, (C) k_C_=k_A_, K_C_=K_A_/4. J_V_ is the total sap flux. k is in 10^−9^ m. s^−1^. MPa^−1^, K in 10^−9^ m^4^. s^−1^. MPa^−1^ and lengths in m.

**Figure 5:**
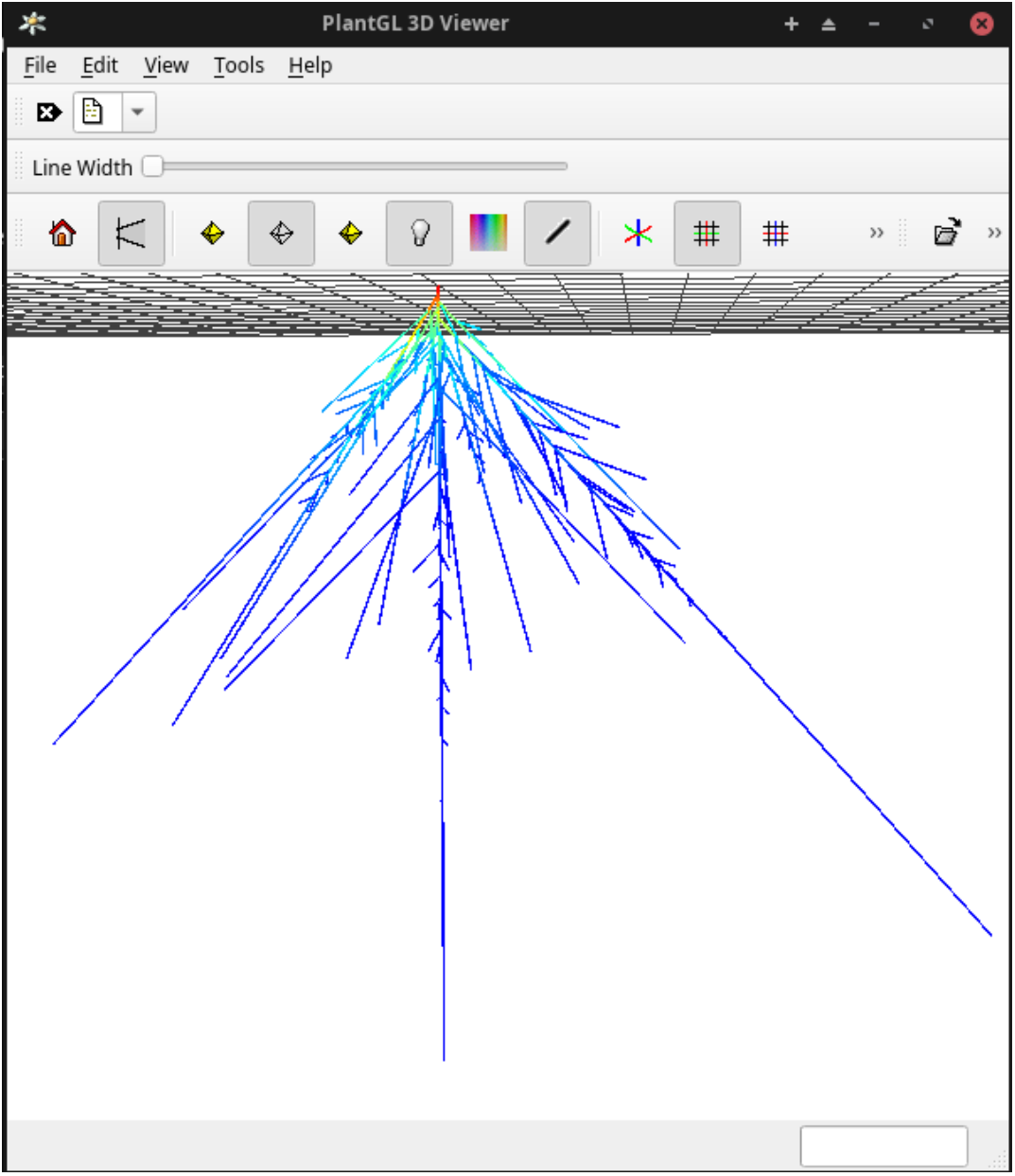
Interactive 3D Plantgl Viewer. Heatmap represents hydrostatic pressure in xylem vessels.

The effect of the radial conductivity value on the root hydraulic architecture can be tested by increasing its value ten-fold (Figure 4 B), by running:

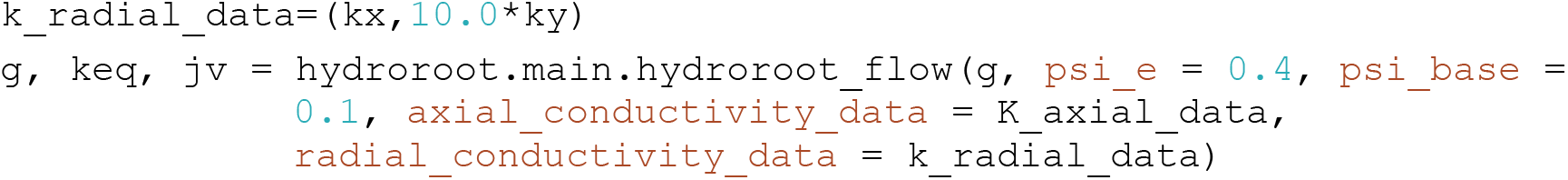
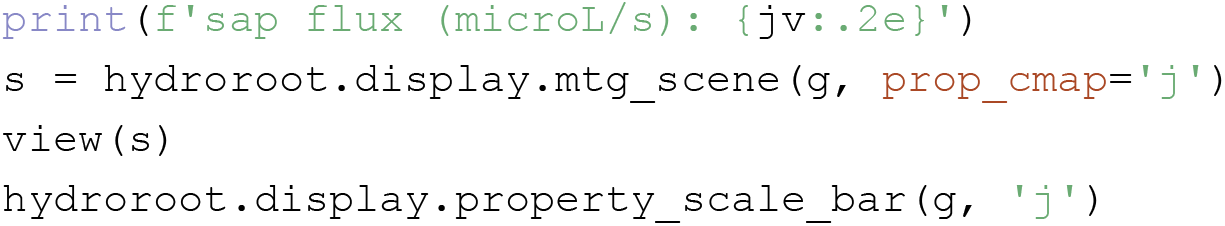

Figure 4 B shows that increasing k makes it less limiting and therefore K becomes more limiting. As a consequence, water uptake follows axial conductance profiles.

We can check the influence of K reduction (Figure 4 C) as follows: setting back k and dividing K, by 4 for example:

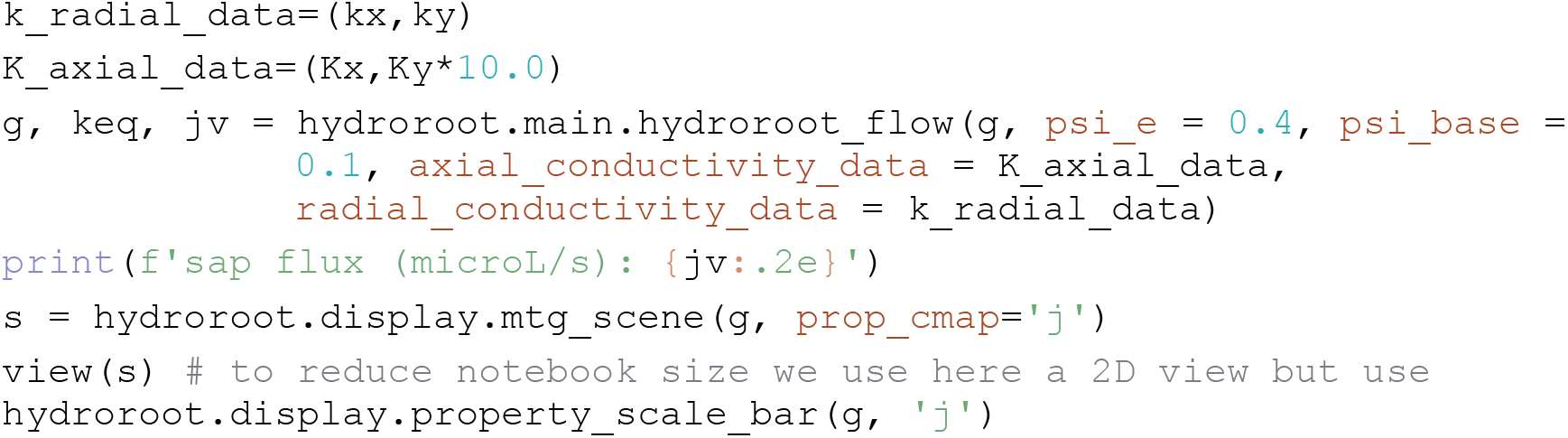

In comparison to Figure 4 A, Figure 4C shows water uptake decrease.

#### 3D viewer

Results can also be displayed in 3D using Plantgl Viewer (Pradal *et al*., 2009):

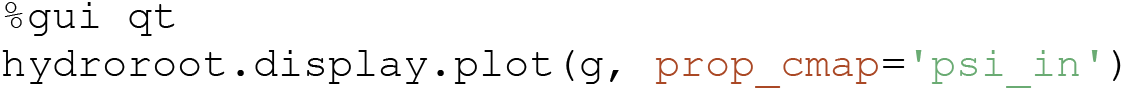

### 4.5. Tutorial 2: simulate hydraulic and solute transport on a maize root

The purpose of this tutorial is to run simulations on a maize root, grown in hydroponic condition using the “solute solver”, and to see how the presence of solutes allows to explain sap flux even without any pressure applied.

#### RSML to MTG

Read the rsml file and convert it to a continuous MTG; this file is located on https://github.com/openalea/hydroroot/blob/main/tutorial/data/maize.rsml.

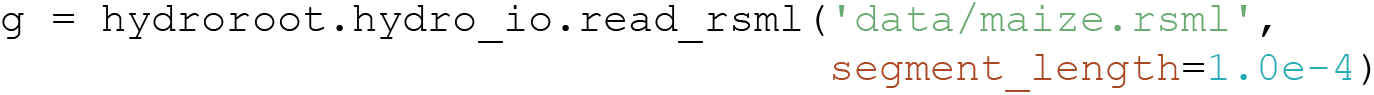

#### Set the root radii

The root radius is set according to the radius of the primary root *r*_*ref*_ = 0.5 10^−3^*m* and the order of the lateral as follows: *r*_*lat*_ = *β*^*order*^*r*_*ref*_. *β* is the radius decreasing factor between root orders. According to Bauget et al (2023) *r*_*ref*_ = 0.5 10^−3^*m* and *β* = 0.4 for maize primary root in standard conditions:

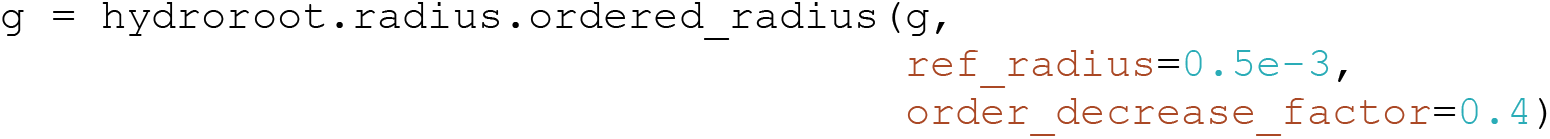

Compute the position of each segment relative to the axis bearing it.

~~~
          g = hydroroot.radius.compute_relative_position(g)
~~~

#### Set hydraulic properties

We set the radial and axial conductivities as profiles, i.e. the property versus the distance to root tip in *m*. The radial conductivity k is in 10^−9^ *m. s*^−1^. *MPa*^−1^ and the axial conductance K is in 10^−9^ *m*^4^. *s*^−1^. *MPa*^−1^

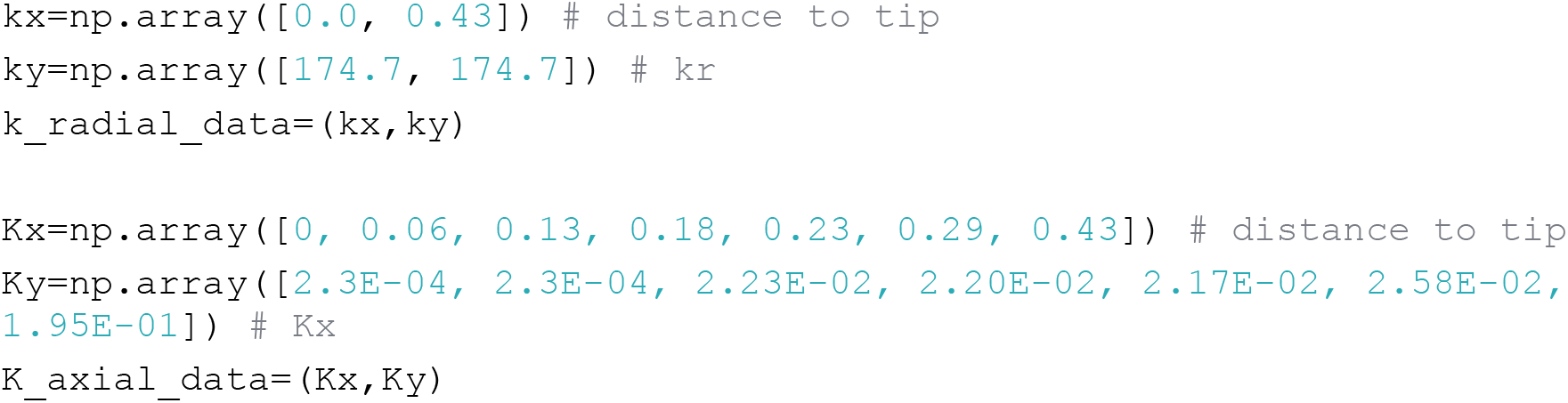

#### Flux calculation without hydraulic potential difference

The simulation is done with the hydrostatic solver for a detopped primary maize root in a hydroponic medium at 0.1 MPa with its base at 0.1 MPa.

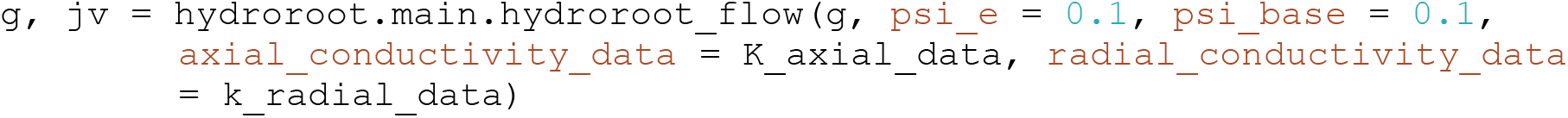

As expected with the hydrostatic solver, there is no outgoing sap flux in the absence of water potential difference.

#### Flux Calculation with solute transport

We now add solute transport using the solute solver with the active pumping rate *J*^∗^ = 1.7 10^−7^ *mol. m*^2^. *s*^−1^, the tissue permeability *P*_s_ = 0.6 10^−9^ *m. s*^−1^ and a solute concentration *C*_*se*_ = 13.69 10^−9^ *mol. m*^−3^ (concentration of the hydroponic solution in Bauget et al. 2023):

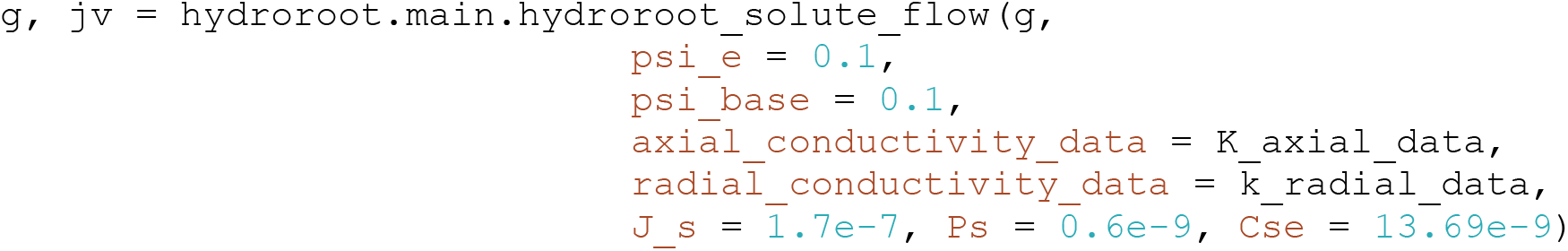

Due to the presence of solutes and an osmotic potential gradient between the external medium and the xylem, the following outgoing sap flux is observed, *J*_*v*_ = 7.7 10^−3^ *μl. s*^−1^ (Figure 6).

**Figure 6:**
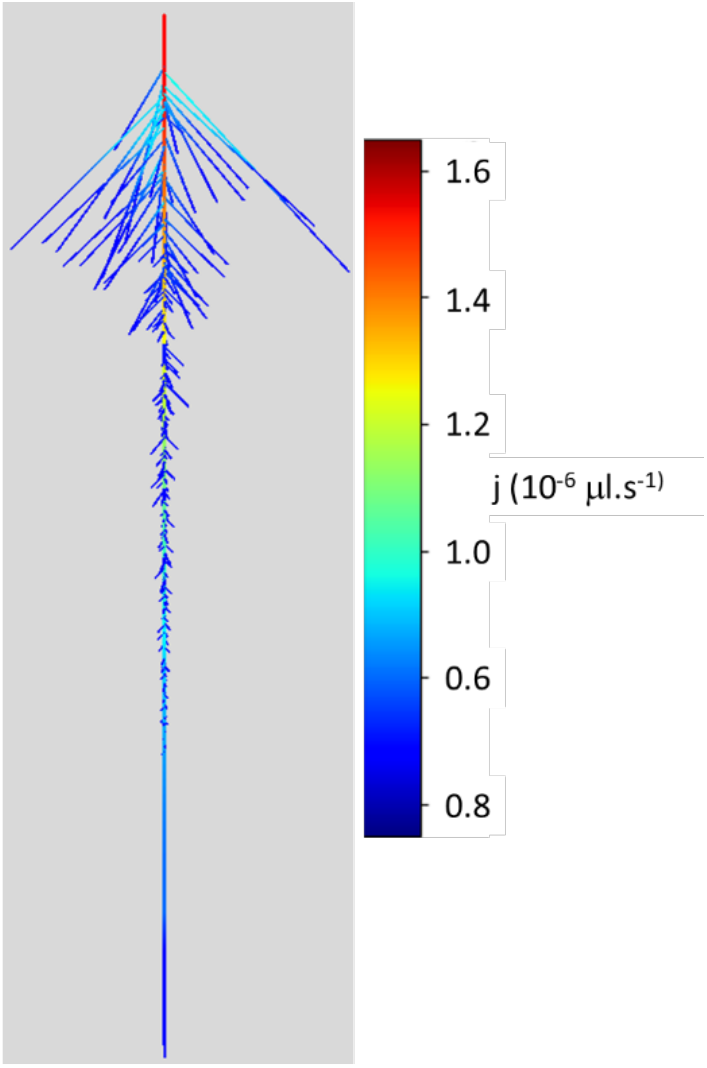
heatmap of water uptake of a maize primary root. Hydoroot solute solver was used without hydrostatic pressure difference. The radial flux is due to osmotic potential gradient between the external medium and xylem vessels.

If, for instance, we increase *J*^∗^ to 1.7 10^−6^ *mol. m*^2^. *s*^−1^, sap flux increases to *J*_*v*_ = 3.1 10^−2^ *μl. s*^−1^.

### 4.6. Tutorial 3: simulate hydraulic on two pearl millet genotypes with varying soil conditions

Two root architectures are simulated for two inbred lines 57 and 109. The root system of pearl millet has three different root types: primary, crown and lateral root types.

#### Modules import

We first import python modules; some are specific to the HydroRoot pearl millet model.

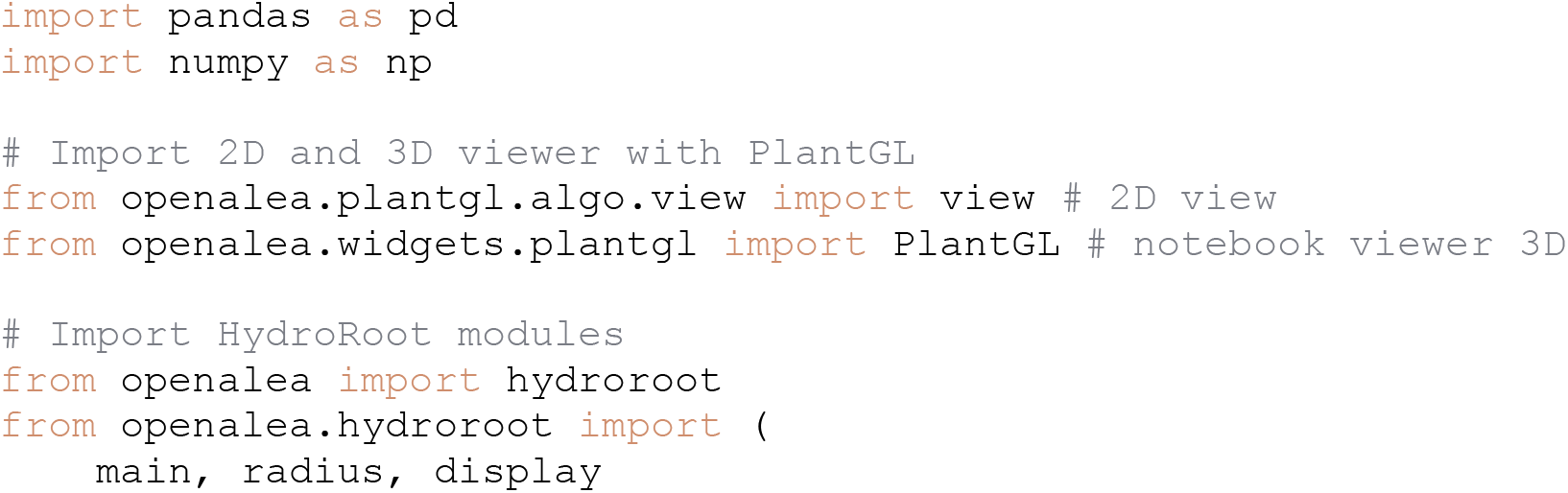
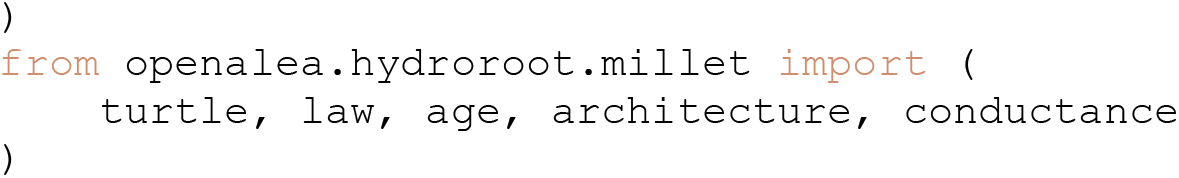

#### Compute RSA

The RSA are generated following architectural parameters given in (Passot *et al*., 2016; Passot, 2016) unless the number of crown roots, which is fixed to 1 and 4 for genotypes 57 and 109, respectively. Lateral lengths are fixed.

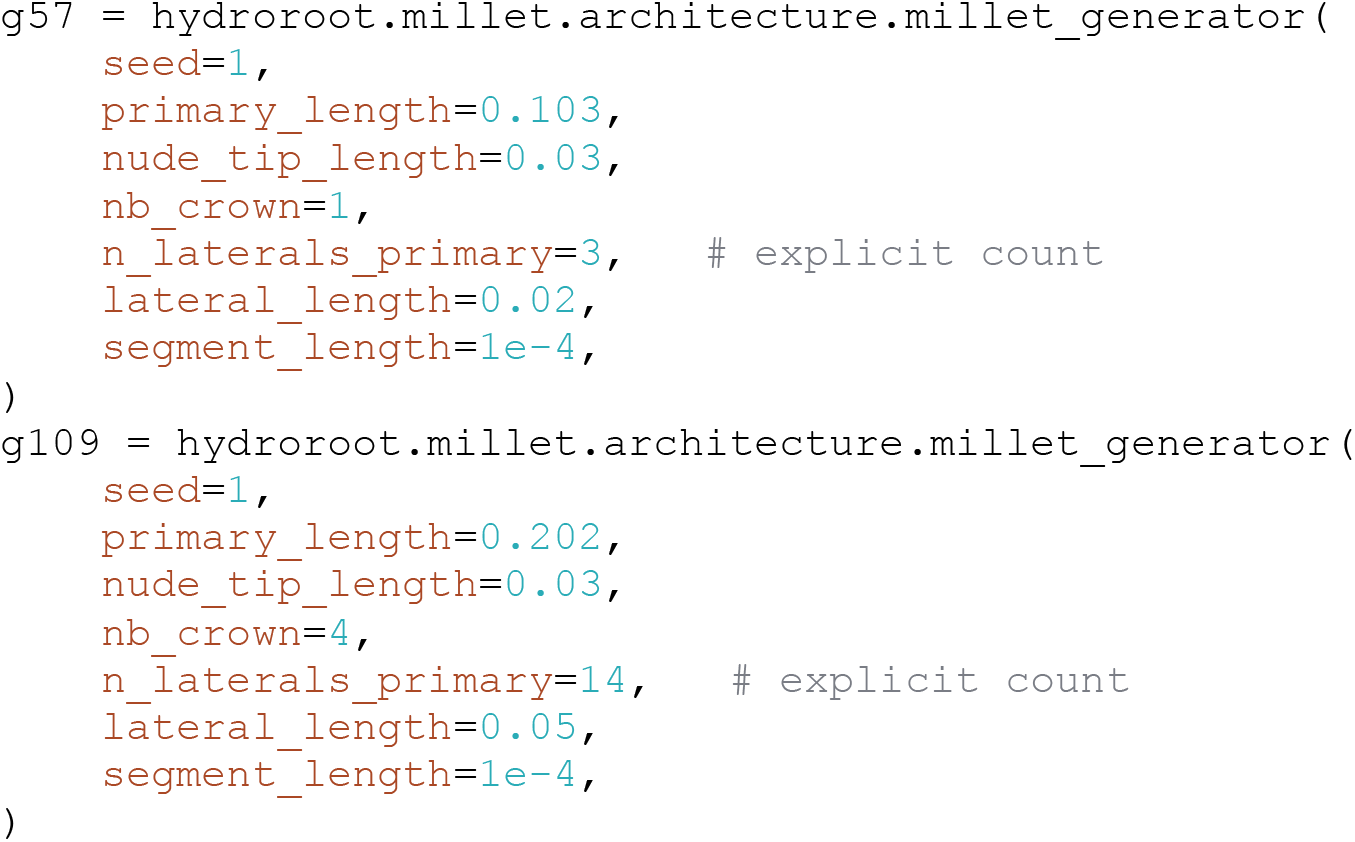

#### Compute length and position

The MTG properties (lengths, position and relative position) are computed.

~~~
          g57 = hydroroot.radius.compute_length(g57)
          g57 = hydroroot.radius.compute_relative_position(g57)
          g109 = hydroroot.radius.compute_length(g109)
          g109 = hydroroot.radius.compute_relative_position(g109)
~~~

#### Compute diameter property using evolution laws

Diameter data are (x,y) csv with position x in cm and root diameter y in µm

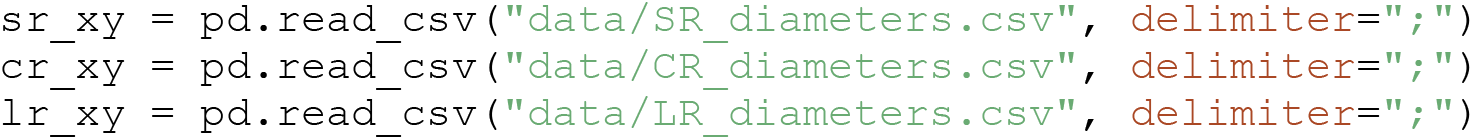

These laws are then fitted with a linear function. Position and diameters are converted to meter, and data are normalized on x and are fitted with an affine function.

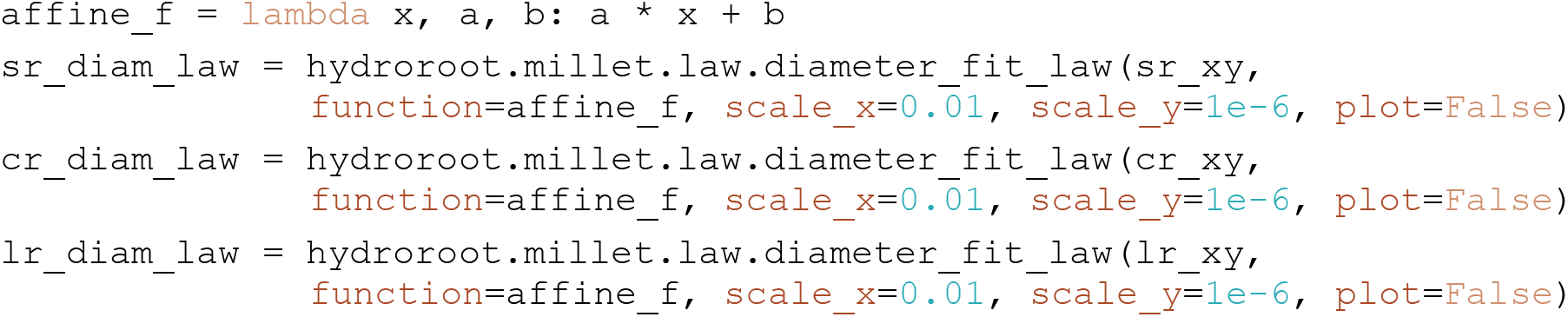

Calculate the MTG diameter property on each vertex according to these laws.

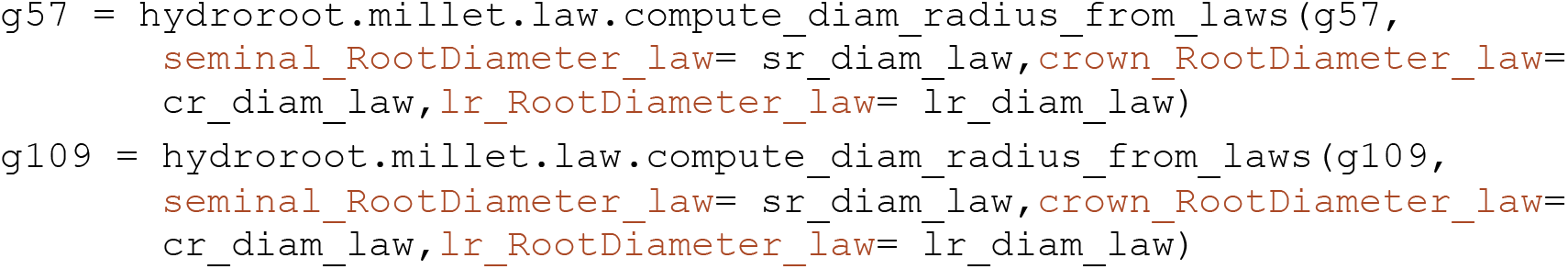

#### Computing Days After Germination (DAG) of root segments

~~~
          g57 = hydroroot.millet.age.compute_dag_with_constant_growth_speed(g57)
          g109 = hydroroot.millet.age.compute_dag_with_constant_growth_speed(g109)
~~~

To visualize age of vertices, we can do for both genotypes (Figure 7):

**Figure 7:**
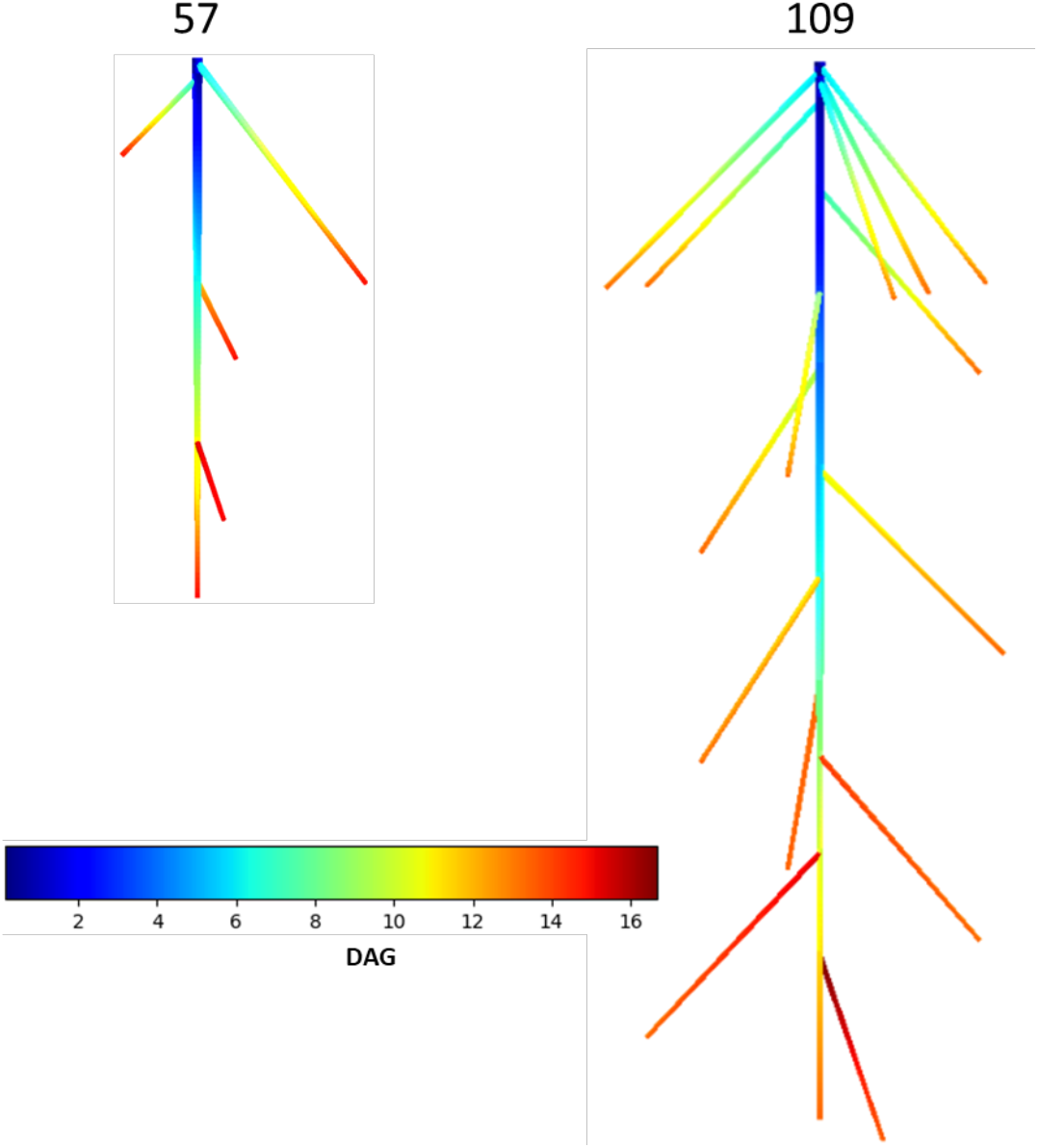
Days after germination (DAG) for two pearl millet genotypes.

**Figure 8:**
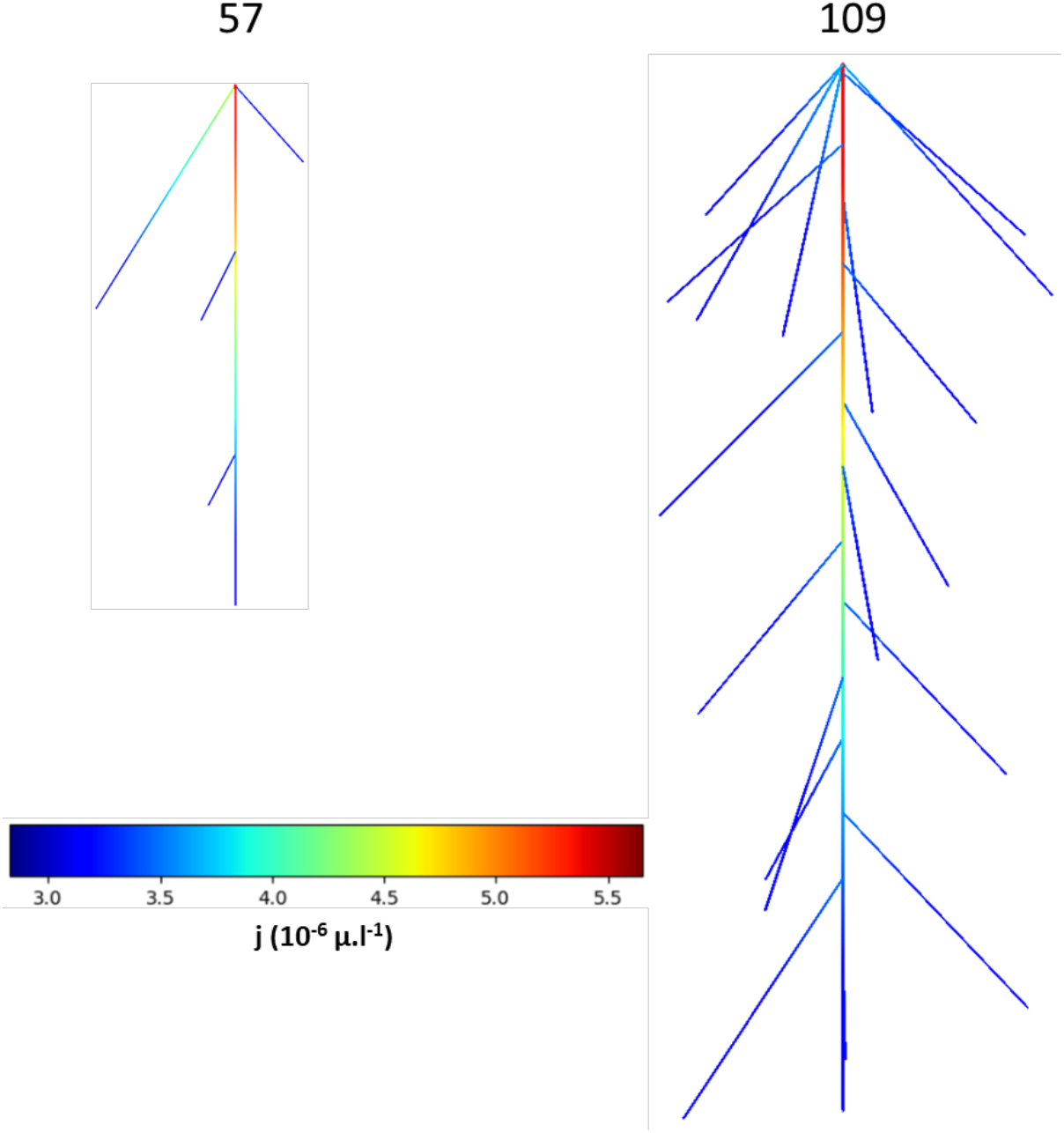
Radial flux j in pearl millet.

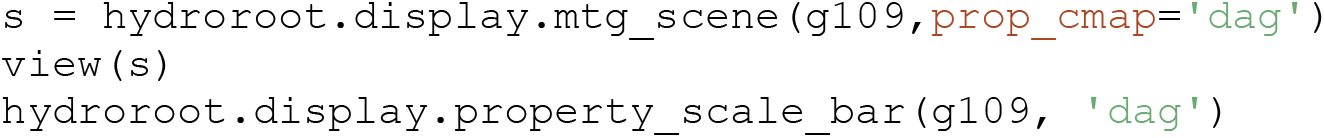

#### Calculation of axial conductances K from the evolution laws

Laws are defined with respect to relative the position x and the unit are 10^−20^ *m*^3^. *s*^−1^. *MPa*^−1^.(Ndour, 2018):

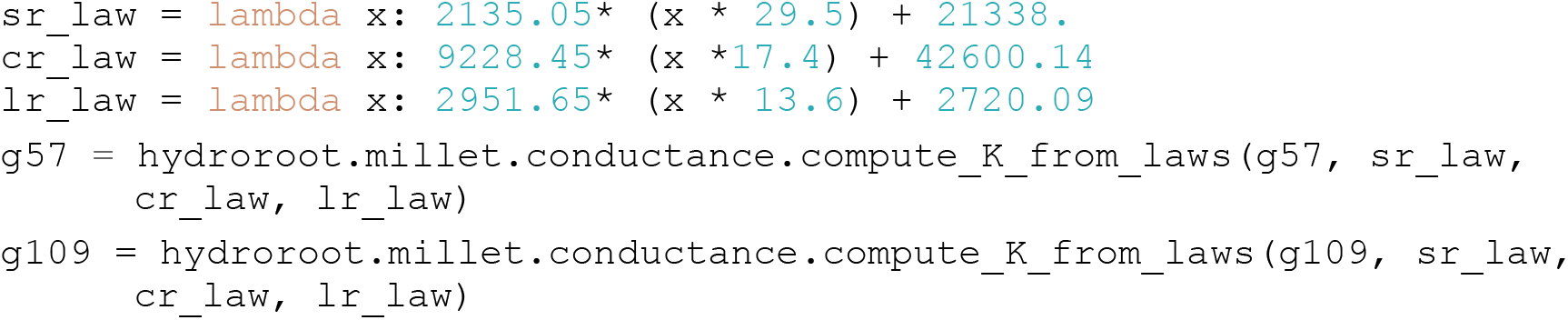

These laws were calculated from xylem vessel geometry using the Hagen-Poiseuille’s law.

#### Compute radial conductances

We set uniform radial conductivities, i.e. the property versus the distance to root tip in *m*. The radial conductivity is in 10^−9^ *m. s*^−1^. *MPa*^−1^ and the same for both genotypes.

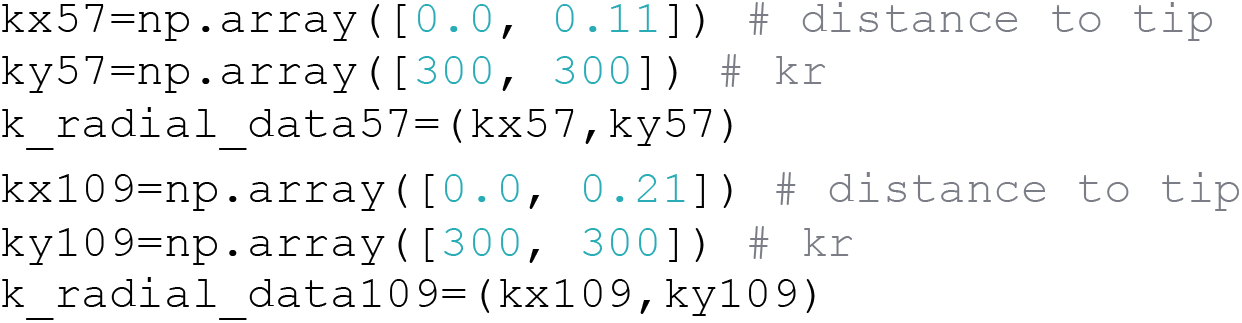

#### Flux calculation

Flux is calculated with the HydroRoot hydrostatic solver.

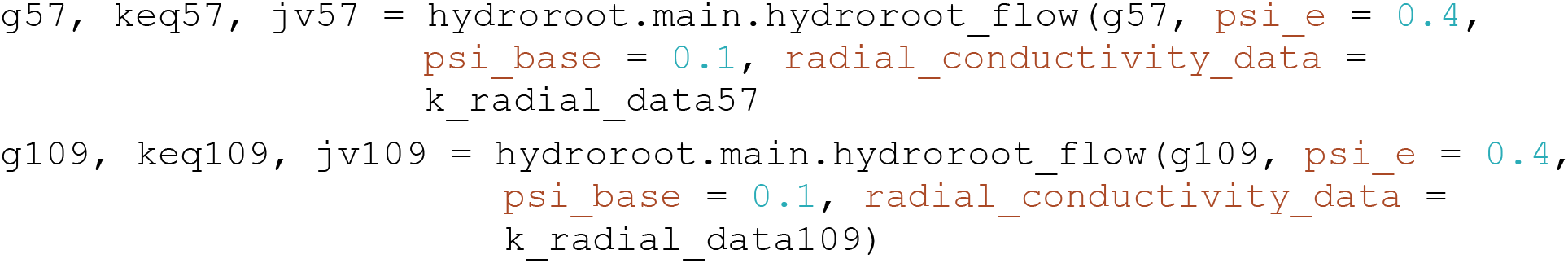

Water uptake can be visualized as follows:

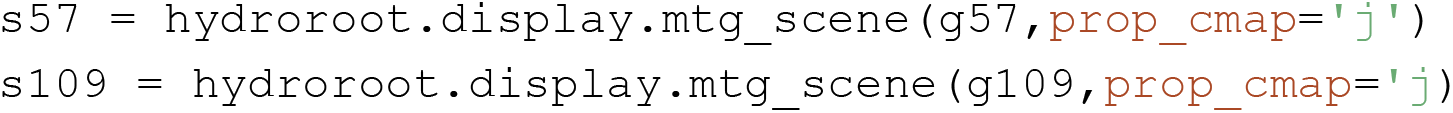

#### Add Soil and compute *Ψ*_*e*_

Hydroroot provides 1D soil whose water potential Ψ_*e*_ varies with depth according to soil type. Simulation with soil implies heterogeneous boundary conditions, unlike simulation in pressure chamber, where conditions are homogeneous.

In this example, two different soils are simulated (Figure 9):

1. A sandy soil where pearl millet traditionally grows in farm conditions.
2. A loamy soil to illustrate effects of the water potential profile.

**Figure 9:**
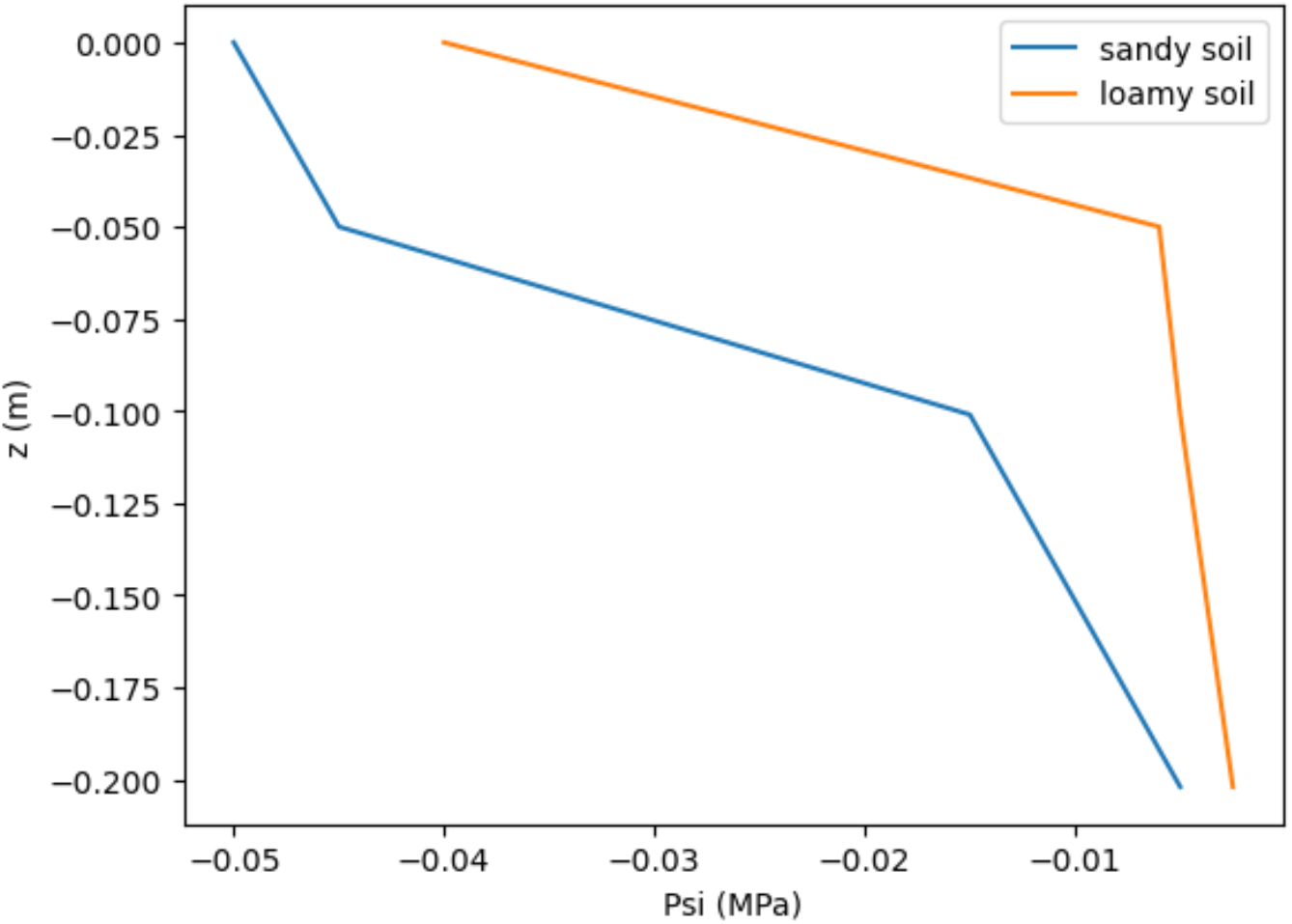
Sandy and loamy soil water potentials.

Soil potential profiles Ψ_*e*_ are negative in MPa versus depth positive in m (Figure 9):

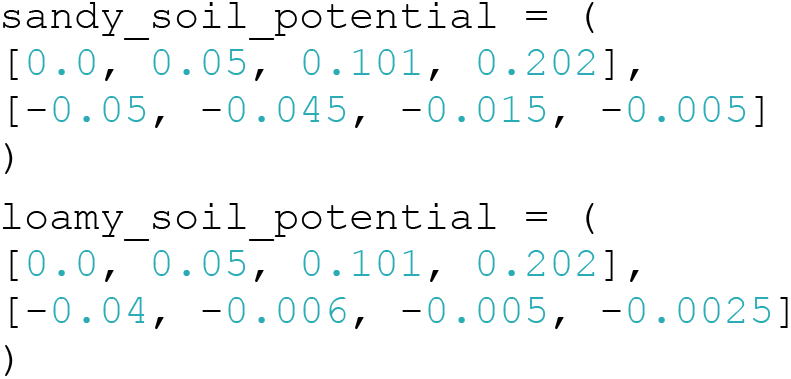

Unlike previous simulation of a root in a pressure chamber, we consider a plant in a soil during day time with transpiration which is simulated with a negative water potential at the collar (MPa):

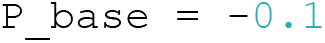

#### xyz position of vertices

The ‘xyz’ MTG property is calculated using a visitor in ‘hydroroot.display.mtg_scene’. Root three dimensional geometries are synthetic, root angles are randomly set according to realistic ranges.

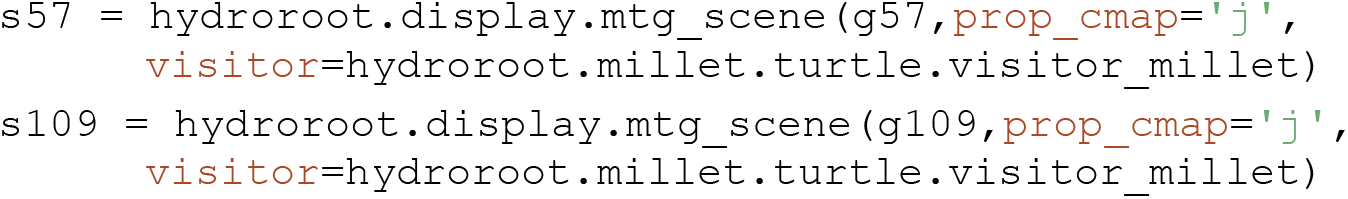

#### Sandy soil simulation

We compute Ψ_*e*_ versus depth and then simulate the flux. We present here the code for genotype 109; codes for genotype 57 are similar.

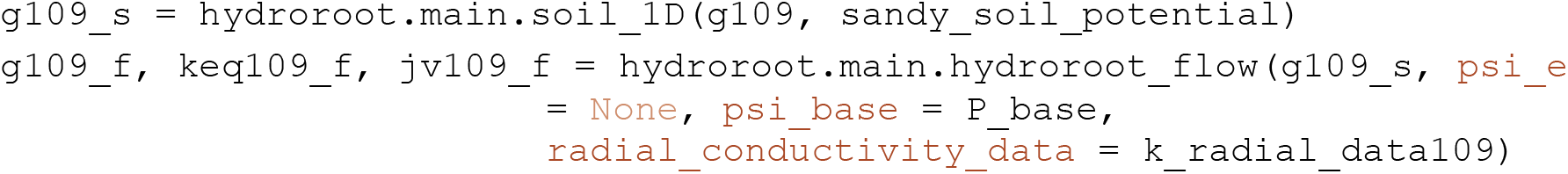

#### Loamy soil simulation

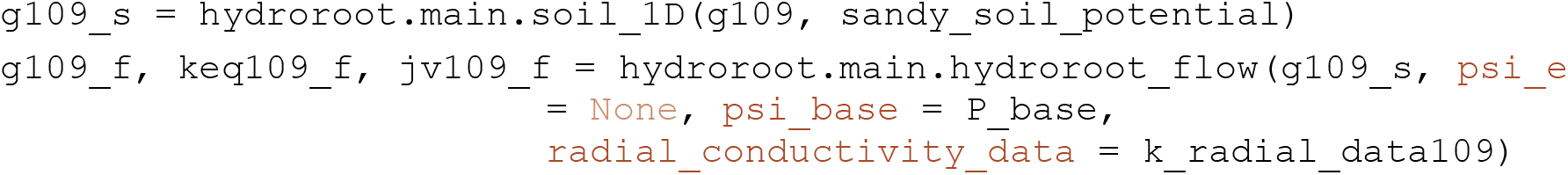

Figure 10 shows the radial flux entering the root for each soil type. As expected, we simulate a higher water uptake in loamy soil. We can also observe the effect of heterogeneous water potential profile with the highest radial flux (in red) localization corresponding to the highest Ψ_*e*_ variation according to depth: between 0.05 and 0.1 m for sandy soil, between 0 and 0.05 m for loamy soil (see Figure 9).

**Figure 10:**
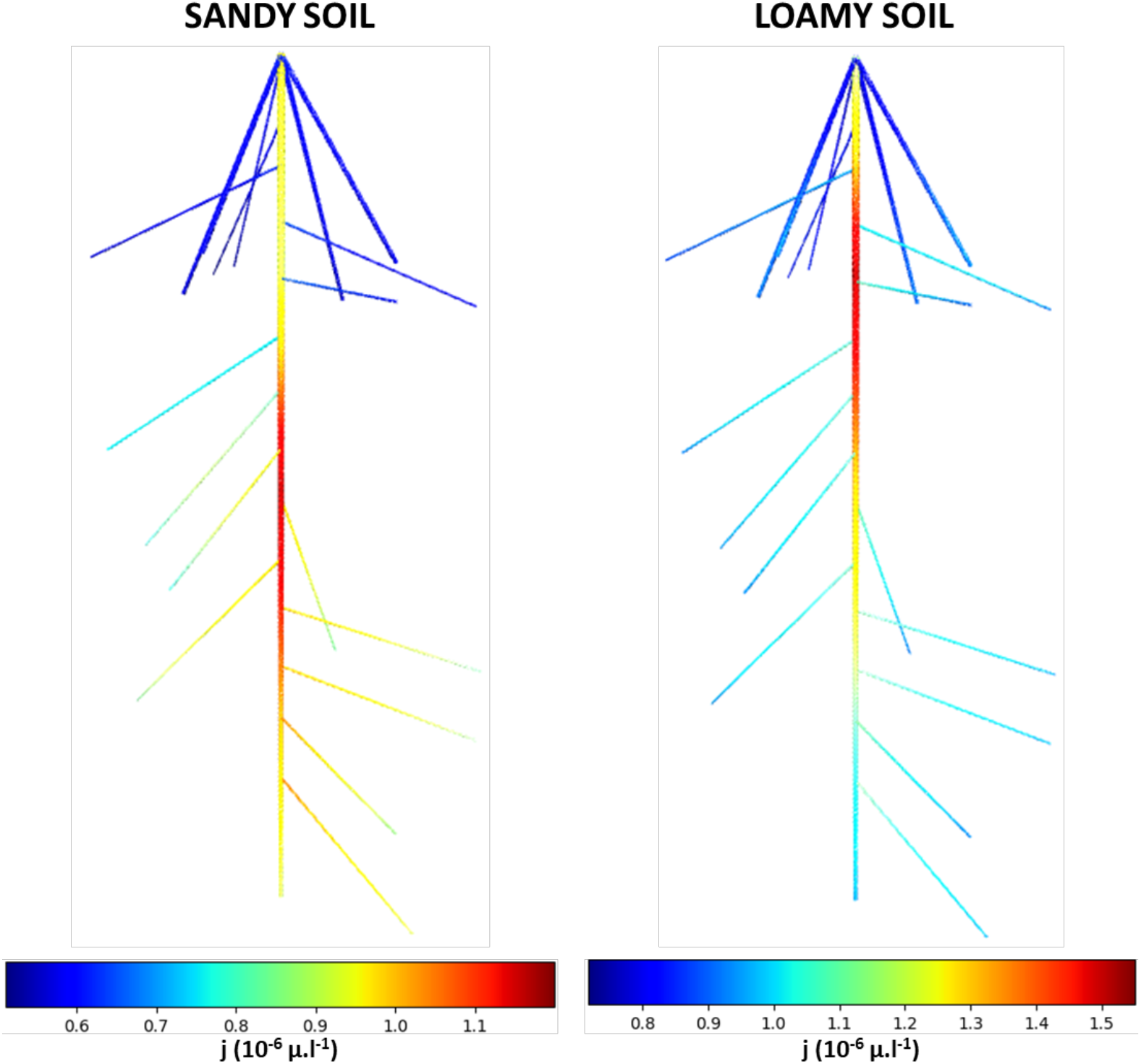
Radial flux of genotype 109 in a sandy and a loamy soil.

### 4.7. Gallery of examples

A gallery of examples is available from the documentation: https://hydroroot.readthedocs.io/en/latest/notebook_examples.html

These examples can also be retrieved from github and interactively executed in Binder:

1. Measured and Simulated architectures: these examples explore the different ways to obtain a RSA, either by generating it or from measurements on real plants.
2. Parameters: a notebook explains how to use the Parameters class of HydroRoot, to group inputs, initialize them, and read a YAML input file.
3. Hydraulic and solute transport parameters adjustments: these tutorials illustrate how to adjust parameters to fit experiments. In particular, one notebook is dedicated to the CnF analysis.
4. Paper Boursiac et al. 2022: a notebook that reproduces all figures and simulations of the paper (Boursiac *et al*., 2022).

## 5. Conclusion

HydroRoot is a water transport modelling framework that have been used 1) to decipher and quantify the role of radial conductance (aquaporins), axial conductance (xylem anatomy) and root system architecture on root water uptake, 2) to phenotype root hydraulic architecture by inverse modelling in well-watered and water-deficit conditions, and 3) to simulate water uptake for various plants (Arabidopsis, maize, pearl millet) with or without soil boundary conditions.

HydroRoot source code is publicly available on Github under open-source license. A documentation with a gallery of examples is available on https://hydroroot.readthedocs.io/.

The integration of HydroRoot in OpenAlea will allow to consider hydraulic architecture with water-mediated Nitrogen uptake like in the OpenAlea module Root-CyNAPS (Gérault *et al*., 2026) and connect root hydraulic architecture with HydroShoot (Albasha *et al*., 2019), a OpenAlea module modeling shoot hydraulic, to build a whole plant FSPM.

